# Biomolecular condensate microstructure couples molecular and mesoscale properties

**DOI:** 10.1101/2025.03.25.645354

**Authors:** Daniel Tan, Dilimulati Aierken, Pablo L. Garcia, Jerelle A. Joseph

**Affiliations:** Department of Chemical and Biological Engineering, Princeton University, Princeton, NJ 08544, USA; Omenn–Darling Bioengineering Institute, Princeton University, Princeton, NJ 08544, USA

## Abstract

Biomolecular condensates, including those formed by prion-like low complexity domains (LCDs) of proteins, are typically maintained by networks of molecular interactions. Such collective interactions give rise to the rich array of material behaviors underlying condensate function. Previous work has uncovered distinct LCD conformations in condensates versus dilute phases, and recently, single-component LCD condensates have been predicted to exhibit microstructures with “small-world” networks—where molecular nodes are highly clustered and connected via short pathlengths. However, a framework linking single-molecule properties, condensate microstructure, and macroscopic material properties remains elusive. Here, we combine molecular simulation and graph-theoretic analysis to reveal how molecular features encode condensate microstructure, which impacts molecule-scale conformations and droplet-scale material properties. Using a residue-resolution coarse-grained model, we probe condensates comprising natural LCD sequences and generalize our findings by varying composition and patterning in binary sequences of hydrophobic and polar residues. We show that non-blocky sequences form condensates with small-world internal networks featuring “hubs”—molecules responsible for global connectivity—and “cliques”, molecular clusters bound by persistent short-ranged associations. Cliques localize near interfaces without a secondary phase transition, suggesting a role in mediating molecular partitioning and condensate aging by tuning interfacial material properties. Moreover, we demonstrate that network smallworldness predicts droplet surface tension. We also track single-molecule structure and dynamics inside condensates, revealing that internal heterogeneity at the single-molecule level is systematically encoded by network topology. Collectively, our work establishes multiscale structure–property relationships in LCD condensates, providing general principles for designing and interpreting condensates with complex internal organization and material properties.

---

Biomolecular condensates are membraneless organelles inside living cells that exhibit a wide range of material behaviors and functions. Phase separation is a leading mechanism that accounts for condensate formation [1–4]. In this framework, proteins and nucleic acids minimize free energy by demixing from the cytosol or nucleoplasm, leading to two or more distinct liquid phases [5, 6]. Unlike simple liquids, evidence suggests that diverse sets of specific and nonspecific interactions between biomolecules yield condensates with nontrivial internal architectures—i.e., transient percolated networks [7–12]. This network microstructure is thought to relate to molecular conformations in the dense phase, as well as to the material properties of condensates [11, 13–15]. Many experiments and simulations have revealed how molecular properties change in condensates compared to dilute phases [10, 16–22]. Recent experiments have also demonstrated that condensates display a diverse array of viscoelastic behaviors [11, 23–30], and further, that condensate viscoelasticity evolves over time—leading to dynamically arrested states associated with pathological condensate aging [28, 31–33]. While the complex features of condensates have been observed at both single-molecule and droplet scales, we lack a fundamental understanding of the principles that connect molecular-level interactions and conformations to macroscopic material properties and functions. A key missing element in relating phenomena across these length scales is a systematic characterization of the condensate microstructure that accounts for complex internal organization and emergent material properties.

Intrinsically disordered regions (IDRs) are among the key components of proteins involved in intracellular phase separation and condensate formation [34]. Prion-like lowcomplexity domains (LCDs) are exemplary instances of IDRs in biomolecular condensation: LCD sequences contain strongly interacting “sticker” residues that drive clustering and phase separation, as well as “spacer” residues interspersed between stickers that modulate solubility and intermolecular interaction strengths [13, 15, 35–38]. These sequence architectures enable the formation of complex networks of reversible physical crosslinks underlying condensates [9, 39]. Recent experimental advances and simulation approaches have begun to observe the rich internal organization and heterogeneities associated with condensate microstructures [10, 12, 40]. Specifically, Farag et al. first l everaged l attice s imulation a nd g raphical network analysis to predict the inhomogeneous connectivity of networks underlying LCD condensates, noting a “smallworld” graph structure globally connected by a small subset of highly connective “hubs” [10]. These results were recently supported by experimental studies revealing the inhomogeneous, network-like internal organization of single-component LCD condensates [12]. Similar network analyses to Ref. 10 have been employed to probe molecular networks in two-component condensates [11], to determine the effect of temperature, length, and residue composition on networks underlying multicomponent condensates [41], and to study the effects of sequence patterning and binding site affinity on percolation and phase separation [18]. Despite experimental and computational characterization, the principles that govern how single-molecule sequence and structure give rise to condensate microstructures—and how microstructures, in turn, encode material properties—remain poorly defined.

Here, we address this gap by combining molecular dynamics simulation and graph-theoretic analysis to characterize the microstructures of LCD condensates. Importantly, we show how condensate microstructure governs droplet material properties, interfacial features, and single-molecule conformations. We systematically study the interaction networks underlying LCD condensates using a chemically-specific residue-resolution coarse-grained model, Mpipi [42]. We generalize our findings by designing binary sequences composed of tyrosine (Y) and serine (S) residues to investigate the impact of sequence composition and patterning on network topology. We consistently find that LCDs and non-blocky binary sequences form condensates with smallworld microstructures marked by high local clustering of molecules and short global pathlengths. Surprisingly, we discover that graph-theoretic parameters describing the extent of small-world organization are directly proportional to droplet surface tension. We further reveal that biomolecules possess two distinct regimes of interactivity in the smallworld network. One regime is marked by high global connectivity and expanded conformations (“hubs”), while the other is marked by elevated local crosslinking (“cliques”). By quantifying the spatial and temporal dynamics of the condensate microstructure, we find that cliques display confined local movements and exhibit long lifetimes compared to hubs, whose molecular identities are found to be highly transient. In agreement with previous experimental studies of mesoscale inhomogeneities in condensates, we find that these nanoscale clique clusters consistently form near interfaces without a secondary phase transition, suggesting roles in mediating selective molecular partitioning and condensate aging [31, 32, 40].

Our work also demonstrates that the condensate microstructure is shaped by a heterogeneous ensemble of single-molecule conformations in the dense phase. Specifically, we predict power-law-like relationships between network connectivity and single-molecule conformational characteristics including radius of gyration and polymer shape anisotropy, indicating that condensate microstructure can be read out with single-molecule features and vice versa. By systematically varying sequence composition and patterning using the binary sequence model, we show that the small-world internal structure is not achieved by blocky sequences, which tend to form micelles instead of a distinct liquid phase. However, the relationships between molecular behavior and microstructure are conserved in all phase-separated condensates, spanning a wide range of sequence compositions. Taken together, our work establishes multiscale structure–property relationships of LCD condensates—linking molecule-scale behavior to droplet-scale material properties—-and provides a conceptual framework for decoding and engineering condensates with complex internal architectures, material behaviors, and functions.

## RESULTS

### Characterization of phase behavior and material properties of LCD and YS condensates

To investigate the phase and material properties of biomolecular condensates composed of LCD-like molecules, we perform molecular dynamics (MD) simulations of single-component LCD condensates (left panel in Fig. 2a). Here, we adopt Mpipi [42]—a chemically specific residue-resolution model that has been shown to describe well the phase behavior of disordered proteins. In addition to characterizing natural LCD sequences (e.g., FUS-LCD), we design and simulate ten “YS variant” sequences composed of tyrosine (Y) stickers and serine (S) spacers with varying hydrophobicity and blockiness [36, 43]. Using the YS sequences, we systematically vary the fraction and position of tyrosine residues in the polymers to examine how well our findings for naturally-occurring LCDs extend to sticker– spacer-like polymer architectures (Fig. 1b,c).

**FIG. 1.**
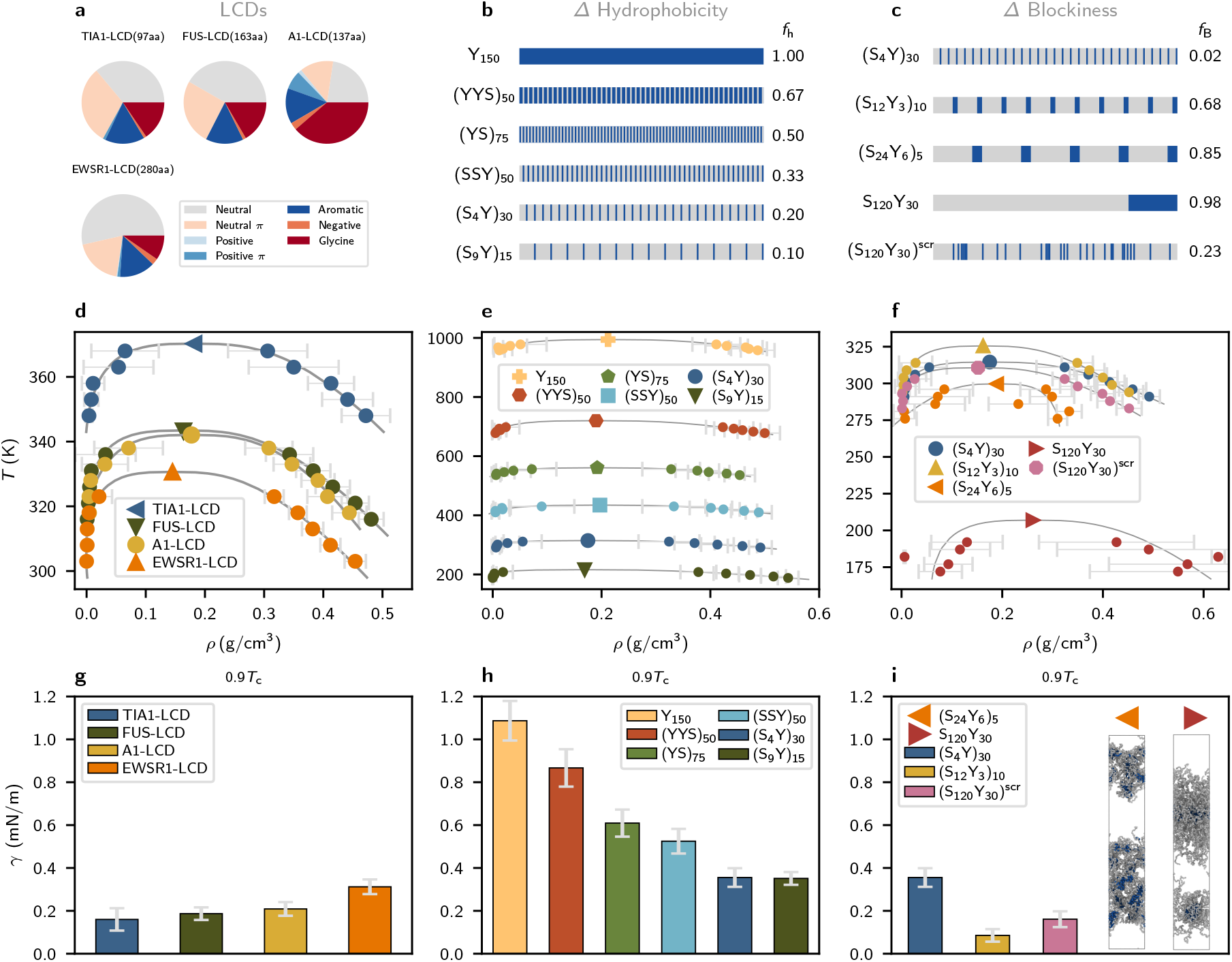
Characterization of phase behavior and material properties of LCD and YS variant sequences. **a** Sequence lengths and compositions of studied LCD sequences. **b** The sequence features of varying hydrophobicity *f*_h_ for YS variant sequences. **c** The sequence features of varying blockiness *f*_B_ for YS variant sequences with fixed hydrophobicity *f*_h_ = 0.20. **d** Phase diagrams for LCD sequences. **e,f** Phase diagrams for simulated YS variant sequences. **g** Computed surface tensions for simulated LCD sequences at 0.9*T*_c_. **h,i** Computed surface tensions for simulated YS variant sequences at 0.9*T*_c_.

We survey biologically relevant condensates by simulating four LCDs known to phase separate under physiological conditions, namely TIA1-LCD, FUS-LCD, hnRNPA1-LCD and EWSR1-LCD. LCD sequence features are shown graphically in Fig. 1(a) and exact sequences are given in the Methods. The critical temperatures *T*_c_ of each sequence are estimated using direct coexistence simulations and the data is fitted using the law of coexisting densities and rectilinear diameters [44, 45], The corresponding phase diagrams are shown in Fig. 1(d). Despite being the shortest sequence (*n* = 97 aa), TIA1-LCD is observed to have the highest critical temperature (*T*_c_ = 370 K) of all LCDs, while the longest sequence EWSR1-LCD (*n* = 280 aa) has the lowest predicted critical temperature (*T*_c_ = 327 K). Notably, TIA1-LCD has the highest fraction of π and aromatic residues (e.g., tyrosine); EWSR1-LCD has the lowest fraction of these residues and the highest fraction of neutral and glycine residues. Consistent with previous reports, we observe that strong π–π and cation–π interactions play outsized roles in driving macromolecular phase separation [36, 39, 46, 47].

All YS sequence variants are constructed with a length of *n* = 150 aa, similar to the average length of the chosen LCD sequences. To study the effect of sequence composition, we varied sequence hydrophobicity (fraction of tyrosine residues, *f*_h_) from *f*_h_ = 0.10 to *f*_h_ = 1.00 over six sequence variants, preserving the near uniform distribution of sticker residues noted for phase-separating prion-like domains [36]: (S_9_Y)_15_, (S_4_Y)_30_, (SSY)_50_, (YS)_75_, (YYS)_50_, and Y_150_. Sequence features are shown graphically in Fig. 1(b), and corresponding phase diagrams are shown in Fig. 1(e). As expected, we predict systematically higher critical solution temperature as hydrophobicity increases. All simulated LCDs have a fraction of aromatic (“sticker”) residues *f*_h_ ≈ 0.14, and the range of critical temperatures observed of LCDs falls roughly within the range of *T*_c_ measured for the YS variants with *f*_h_ = 0.1 and *f*_h_ = 0.2.

To probe the effect of sequence patterning, we design three additional YS variants at a hydrophobic fraction *f*_h_ = 0.20, as the corresponding uniform sequence (S_4_Y)_30_ displayed the closest phase behavior to the LCDs with *f*_h_ = 0.14. We then alter the blockiness of the sequences: (S_12_Y_3_)_10_, (S_24_Y_6_)_5_, and S_120_Y_30_. In addition, we generate a randomly scrambled sequence, (S_120_Y_30_)^scr^, with the same composition. Graphical representations of these sequences are shown in Fig. 1(c) along with their measured blockiness *f*_B_ (see Methods), and corresponding phase diagrams are shown in Fig. 1(f). The critical temperatures and phase boundaries of these patterning variants are all similar to those of LCDs except for those corresponding to (S_24_Y_6_)_5_ and S_120_Y_30_, the two blockiest sequences. These latter sequences form micelles instead of phase-separated condensates.

In addition to characterizing the phase behavior of the sequences, we investigate how molecular sequence affects condensate material properties by computing droplet surface tension. We measure the surface tension of each condensate via direct-coexistence simulations in the slab geometry [48] at 0.9 *T*_c_, where *T*_c_ is the critical solution temperature. We find that surface tension increases with LCD sequence length [Fig. 1(g)]. Longer chains likely enhance surface tension due to confinement and entanglement. On the other hand, condensates formed by uniformly patterned YS sequences show that higher sequence hydrophobicity is proportional to surface tension [Fig. 1(h)], suggesting the dominant role of hydrophobicity in modulating material proper-ties. The effect of hydrophobicity is likely obscured in LCD condensates due to electrostatic repulsion arising from complex sequence features, varying sequence lengths, and only minor variations in LCD hydrophobicity. The patterning variants shown in Fig. 1(i) demonstrate that surface tension decreases with increased sequence blockiness, proceeding in order from (S_4_Y)_30_ to (S_120_Y_30_)^scr^ and (S_12_Y_3_)_10_ (see Fig. 1(c) for sequence blockiness *f*_B_). However, the formation of micelle structures, shown in Fig. 1(i), prevents us from properly calculating the surface tension for (S_24_Y_6_)_5_ and S_120_Y_30_. While the YS model enables a controlled study of sequence effects, the LCD trends suggest that the emergent material properties of the condensate are shaped by an interplay between sequence length, hydrophobicity, and charge patterning. Together, these results point to nontrivial but interpretable relations between sequence composition, condensate phase behavior, and macroscopic material properties.

### Small-world connectivity in LCD and YS condensates predicts surface tension

To properly investigate the microstructure underlying condensates, we simulate each sequence in an isotropic box in the *NV T* ensemble at *T* = 0.9*T*_c_. The isotropic simulation cell permits the formation of spherical droplets at equilibrium, enabling graph-theoretic analyses on networks of finite size. In network representations of condensate microstructures, individual molecules are represented as single nodes, and interacting nodes are connected with unweighted and undirected edges (right panel in Fig. 2a). Specifically, we assign edges between two interacting molecules (i.e., LCDs or YS sequences) *A* and *B* when the interaction potential energy falls below a threshold of − 5*k*_B_*T*. The scalar multiple of *k*_B_*T* tunes the strength of the energetic criterion to unveil more transient (lower multiples) or more long-lived (higher multiples) networks representing the condensate microstructure. In the Mpipi model, the interaction strength encoding the strong hydrophobic Y–Y interaction is 0.42 kcal/mol, which is slightly smaller than the thermal energy *k*_B_*T* = 0.59 kcal/mol at 300 K. A typical distribution of intermolecular interaction energies between pairs of A1LCD molecules at 0.90*T*_c_ is shown in Fig. 2b, where 68% of all recorded intermolecular interactions are stronger than the thermal energy *k*_B_*T*, 33% of all interactions are stronger than 5*k*_B_*T*, and only 14% of all interactions are stronger than 10*k*_B_*T*. Considering that the 5*k*_B_*T* threshold represents an interaction energy greater than five instances of strong, hydrophobic Y–Y interactions, the results reported in the text and figures use 5*k*_B_*T* as a reasonable threshold to define relatively stable interactions and ensure that edges in computed networks represent long-lived associations within the microstructure.

**FIG. 2.**
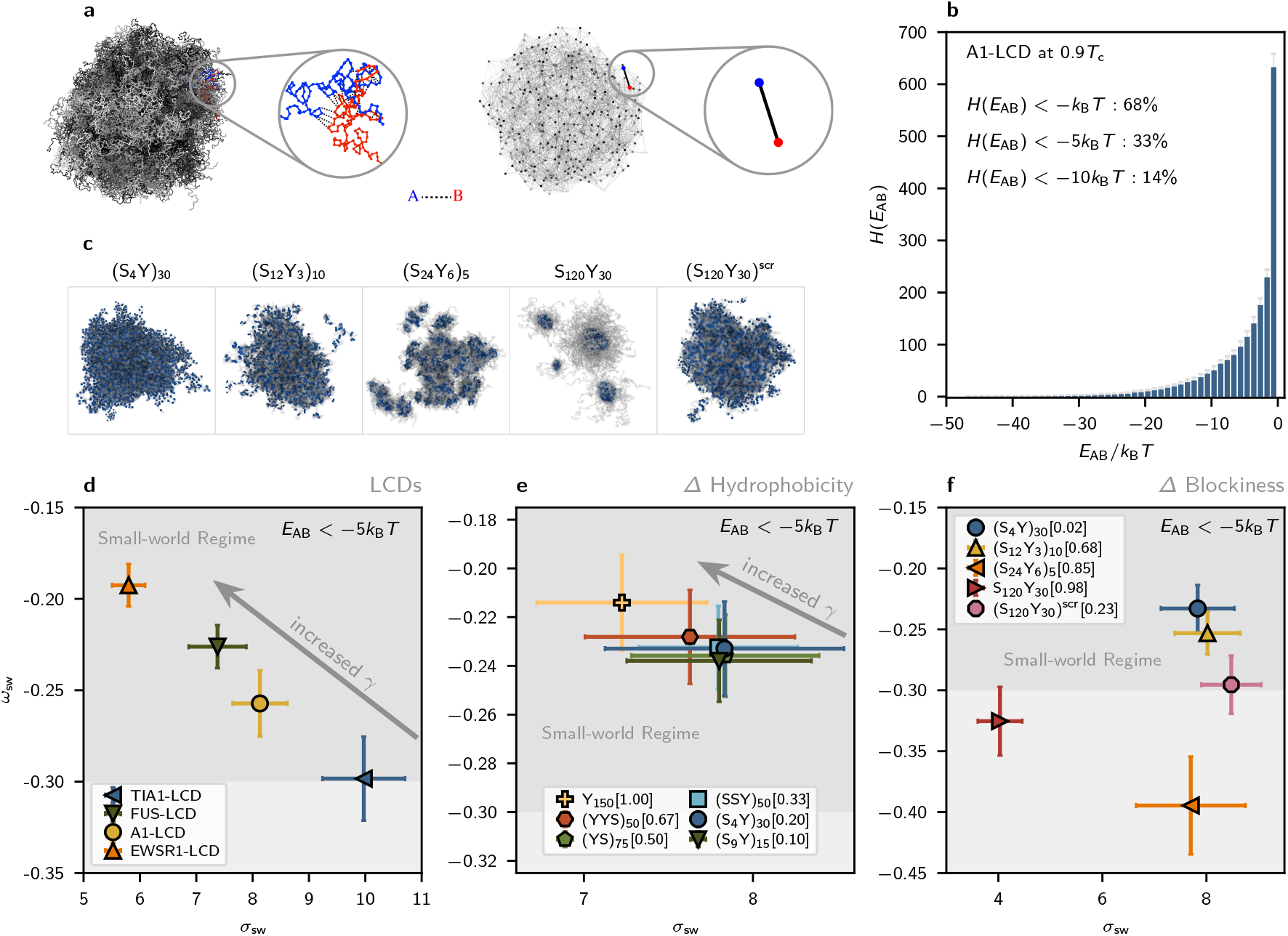
Small-world connectivity in LCD and YS condensates predicts surface tension. **a** Snapshot of a simulated condensate (left) and its corresponding graph representation (right), with two interacting molecules depicted in the insets. In the graph representation, each molecule is taken as a node, and two nodes are connected with an unweighted, undirected edge when the sum of pairwise monomer interaction energies (*E*_AB_) between them exceeds a threshold based on the thermal energy (*k*_B_*T*). All results and figures in this work use the threshold 5*k*_B_*T*. **b** Histogram depicting the distribution of intermolecular contact energies in a simulation of A1-LCD at 0.9 *T*_c_. The 5*k*_B_*T* threshold captures ≈ 33% of all recorded interactions. **c** Morphologies of selected YS variants are shown, with the two blockiest sequences (S_24_Y_6_)_5_ and S_120_Y_30_ showing micellization with core-shell architecture. **d** Small-world parameters *σ*_sw_ and *ω*_sw_ for single-component LCD condensates. An arrow is overlaid to show that droplet surface tension increases as the microstructure approaches ideal small-world networking (*ω*_sw_ towards 0). **e** Small-world parameters *σ*_sw_ and *ω*_sw_ for condensates formed by YS sequences with varying hydrophobicity. As in (d), an arrow is overlaid to show that droplet surface tensions increase as microstructures become more small-world-like (*ω*_sw_ towards 0). **f** Small-world parameters *σ*_sw_ and *ω*_sw_ for condensates formed by YS sequences with *f*_h_ = 0.2 and varying patterning. The two blockiest sequences (S_24_Y_6_)_5_ and S_120_Y_30_ form micelles with underlying networks that are distinctly not small-world-like.

Using this procedure, we find that condensates formed by LCDs and non-blocky YS sequences consistently display small-world network microstructures that span the dense phase [Fig. 2(d,e)]. Small-world networks are formally characterized by high clustering coefficients and low average shortest pathlengths [49, 50]. We quantify network smallworldness using the graph-theoretic estimators *σ*_sw_ and *ω*_sw_, which measure the average clustering coefficient (*C*) and average shortest pathlength (*L*) between arbitrary nodes in the graph [51–54]. The equations used to compute these graph parameters are described in the Methods. Values of 0 *< σ*_sw_ *<* 1 indicate that clustering is low or average shortest pathlengths are long compared to equivalent Erdős– Rényi (ER) random graphs, and *σ*_sw_ ≈ 1 indicates that the network is organized like an ER random graph. Characteristic small-world values *σ*_sw_ *>* 1 come from high clustering coefficients and average shortest pathlengths that are shorter than or comparable to those in ER random graphs. The second estimator *ω*_sw_ is bounded between -1 and 1, where *ω*_sw_ = − 1 corresponds to a regular, lattice-like graph structure and *ω*_sw_ = 1 corresponds to a random-graph structure. The small-world region *ω*_sw_ ≈ 0 describes a graph structure that is both highly clustered—like regular lattices—and has short average path lengths, like ER random graphs [49, 53].

The balance of high clustering and short path lengths underlies the resilience and conduciveness of the small world network to efficient, high-fidelity transfer: most nodes are well-connected to local nodes in clustered “neighborhoods” (graph “cliques”), and these neighborhoods are globally linked through a small subset of highly connected “hub” nodes that act as highways mediating pairwise node relations through shortest paths.

Fig. 2d shows that all LCD condensates exhibit microstructures defined by small-world interaction networks with characteristic *σ*_sw_ *>* 1 and *ω*_sw_ ≈ 0 values. These results are consistent with those from lattice simulations [10], which found that single-component A1-LCD condensates have interaction networks exhibiting small-world topologies. Strikingly, a clear trend emerges between the smallworldness of the microstructure and the condensate surface tension. As condensate microstructure becomes more small world-like with *ω*_sw_ approaching 0, the macroscopic surface tension increases [Fig. 1(g) and overlaid arrow in Fig. 2(d)]. The YS sequences with uniform sticker patterning also display small-world microstructures across the full range of sequence hydrophobicities examined [Fig. 2(e)]. Notably, the qualitative relationship between network small-worldness and droplet surface tension is maintained. With decreasing sequence hydrophobicity, YS polymers form condensates with progressively lower surface tensions and greater deviations from ideal small-world structure [Figs. 1(h) and 2(e)]. The patterning variants shown in Fig. 2(f) suggest that increased sequence blockiness disrupts the small-world connectivity of the microstructure. Indeed, the blockiest sequences (S_24_Y_6_)_5_ and S_120_Y_30_ form micelles instead of phase separating. The effects of heterogeneous interactions are most dominant in these systems, and remarkably, these sequences show the greatest deviation from ideal smallworldness as measured through *ω*_sw_. Together, our results demonstrate how molecular sequence impacts the condensate microstructure and how this microstructure directly relates to the condensate surface tension.

### Molecular hubs and cliques are spatially segregated

To further characterize condensate microstructure using underlying molecular networks, we analyze the spatial distribution of small-world topological features at the singlemolecule level. Small-world networks rely on “hubs”, members of a small subset of highly connective nodes, to lower average pathlengths by mediating many of the shortest paths connecting arbitrary node pairs. Small-world networks also contain “cliques”, locally fully connected neighborhoods of nodes [55]. In the context of biomolecular condensates, cliques are clusters of closely interacting macromolecules. Clique substructures tend to be bridged to other cliques through hubs, which promotes efficient flow and network resilience—crucial properties of small-world topologies that drive their adoption in engineered and natural settings [55].

We thus examine the spatial distributions of hub molecules and clique molecules in dense phases. The typical distributions of hub and clique molecules in the graph representation are depicted in Fig. 3a. Hubs are identified by high betweenness centralities *C*_B_, a measure of the extent to which a single node lies along shortest paths between arbitrary node pairs in the graph. Detailed calculations are shown in the Methods. Here, the top 10 hub molecules (i.e., highest betweenness centrality *C*_B_) are colored in red and the 10 largest cliques are shown in blue. Next, we quantify the spatial distribution of hubs and cliques in the condensates. As shown for simulations of FUS-LCD in Fig. 3b, mass density profiles are roughly uniform within simulated condensates, featuring well-defined interfaces. However, despite homogeneous density profiles, the distributions of hub molecules and clique molecules are distinctly heterogeneous: cliques are located closer to the interface than are hubs, which are distributed throughout the volume of the condensate. We further confirm that such distributions are not a special case for FUS-LCD but are generic features of condensates formed by LCDs and uniformly patterned YS variant sequences, as characterized in Fig. 3d. We also confirm that this effect is not due to finite-size effects by simulating an analogous system composed of 3375 copies of FUS-LCD, shown in Supplementary Figure S5. We find that the observed distribution of hubs and cliques is persistent in larger systems.

**FIG. 3.**
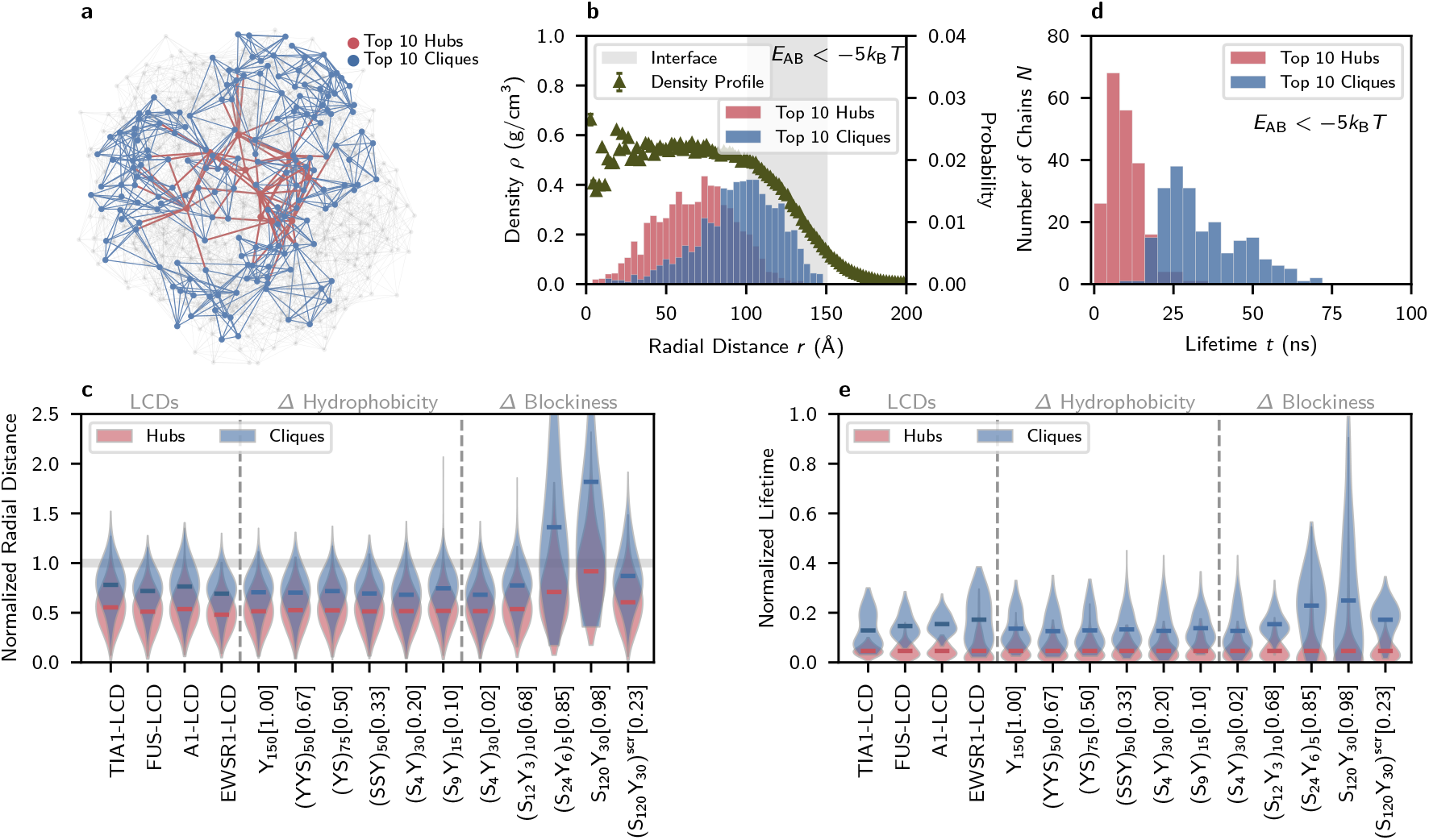
Network hubs and cliques are spatially and temporally distinct. Constructed interaction networks and graph-theoretic analyses use an energetic threshold of 5*k*_B_*T* to define network edges representing long-lived intermolecular associations. **a** Graph representation of an A1-LCD condensate (512 chains). Hubs are colored red and cliques are colored blue. Molecules colored gray are neither hubs nor members of the largest cliques. **b** Spatial distribution of hub molecules and clique molecules within a simulated FUS-LCD condensate (216 chains, *T* = 270 K). The clique molecules are closer to the interface than hub molecules. A radial distance of zero represents the center of mass of the condensate. The density profile is shown as triangles and the interface region is shaded in gray. An analogous profile from a simulation of 3375 FUS-LCD chains at *T* = 300 K is shown in Supplementary Figure S5; the observed distribution of hubs and cliques is persistent in larger systems and at higher temperatures. **c** The spatial distribution of hubs and cliques for all simulated sequences in terms of radial distance, normalized by the distance from each condensate’s center of mass to its interface (overlaid grey bar). **d** The lifetime distribution of hub molecules and clique molecules are shown over 200 nanoseconds for FUS-LCD (216 chains, *T* = 270 K). Hub molecules are transient; the majority of hub molecules remain hubs on relatively short timescales, while molecules within cliques remain members of cliques for substantially longer periods of time. **e** Distributions representing the normalized lifetimes of hub molecules and clique molecules are shown for all simulated sequences.

Additionally, we find that the spatial distributions of hubs and cliques appear to be weakly dependent on sequence patterning. Between (S_4_Y)_30_, (S_120_Y_30_)^scr^, and (S_12_Y_3_)_10_, greater sequence blockiness leads to slightly more pronounced distinctions between the spatial distributions of hubs and cliques [Fig. 3(d)]. The blockiest sequences (S_24_Y_6_)_5_ and S_120_Y_30_ do not phase separate and instead form micelles with core-shell architecture [Fig. 2(c)]. Hubs and cliques identified in micelles are not meaningful organizing features, as molecular packing in the hydrophobic micelle core enables global connectivity and elevated local clustering simultaneously. Due to this, no conclusive relation between sequence blockiness and hub–clique mesoscale inhomogeneity can be determined.

Collectively, our findings reveal that the small-world microstructure of LCD-like condensates predictably encodes mesoscale heterogeneities within dense phases, organizing molecules into distinct spatial regimes of interactivity. Molecules in dense phases act as hubs, contributing to global network connectivity, while molecules near the interface tend to form clique clusters defined by tight local associations. This spatial partitioning of network roles reflects a non-random, emergent organization even in single-component condensates, offering a mechanistic link between network topology and spatial patterning. Notably, our results recapitulate previous experimental observations of nanoscale molecular clustering at the interfaces of multicomponent condensates [40], suggesting that the emergence of spatial inhomogeneities from the microstructure is broadly conserved in phase-separated macromolecular assemblies. These insights emphasize the importance of treating the condensate interface not as a passive boundary but as a functionally and structurally distinct region, whose qualities strongly influence condensate material properties, biological function, and aging behavior.

### Molecular hubs and cliques exhibit distinct lifetimes

Given that networked microstructures lead to pronounced spatial inhomogeneities in condensates, an important open question is whether these microstructures also engender dynamical inhomogeneities, or differences in the temporal stability of network roles. Such inhomogeneities, especially as pertaining to interfacial properties, could have direct implications for condensate material properties. To explore this, we analyze the lifetimes of hubs and cliques within LCD and YS condensates to study the dynamics of the microstructure (see Methods). We measure molecular lifetimes based on their network identities: we quantify the fraction of time that a molecule is identified as a hub or as belonging to a clique over a continuous 200 ns trajectory sample. For FUS-LCD, we find that cliques exhibit significantly longer lifetimes than hubs [Fig. 3(c)]. In fact, for all LCDs probed, individual molecules very scarcely serve as connective hubs for more than 1–2 ns in our simulations, while members of cliques remain in those cliques for substantially greater time fractions [Fig. 3(e)]. This clear temporal separation of hubs and cliques is also observed for non-blocky YS sequences [Fig. 3(e)], and the behavior appears to be conserved across all uniformly patterned YS variants. Despite their longevity, molecular transfer in and out of clique clusters is still visible in Fig. 3(e), namely via the lack of clique molecules that remain in their cliques for any greater than ≈ 40% of the trajectory sample.

YS pattern variants suggest that sequence blockiness is highly related to clique longevity: the blockiest phase-separating sequence (S_12_Y_3_)_10_ displays the clearest distributional distinction between hub and clique lifetimes, and hub and clique distributions overlap more as sequence blockiness decreases [Fig. 3(e)]. Further, LCD condensates show that clique lifetimes are dependent on sequence length and diversity (Fig. 3e)—effects not captured in the binary YS sequences. For example, EWSR1, the longest and least hydrophobic LCD tested, exhibits both longer clique lifetimes and a broader distribution of clique lifetimes compared to other LCDs. TIA, the shortest, most hydrophobic, and least uniformly patterned LCD studied, has a broad and bimodal distribution of clique lifetimes. While the general temporal distinction between hubs and cliques in LCD and YS condensates remains conserved, these results imply that the properties of molecular clusters at interfaces, including their stability and local dynamics, can be fine-tuned with sequence patterning.

Taken together, these results suggest that the condensate microstructure obeys counterintuitive dynamics. The backbone of the small-world network consists of highly associative hub molecules who rapidly interchange roles, while the peripheral networks of local interactions between clique molecules are macroscopically more time-stable. This “decentralization” of hublike connectivity in the microstructure may represent a physical mechanism of resilience against network failure due to aberrant single-chain behavior. Moreover, experiments have shown that nanoscale molecular clustering is linked to reduced molecular diffusion in dense phases [40]. The relative stability of clique clusters, along with their localization to condensate interfaces, suggests that clustering may play a role in selective molecular recruitment and dynamic confinement. These results provide a framework for understanding and interpreting condensate function and mesoscale inhomogeneity as emerging from interaction networks, properties of which can be carefully tuned with sequence composition and patterning.

### Conformations of molecules in biomolecular condensates are dependent on network topology

Recent work has highlighted that LCDs adopt heterogeneous conformational ensembles within condensates, with molecular properties that vary depending on the local microenvironment. In particular, differences in average molecular size (i.e., radius of gyration *R*_g_) have been reported between proteins at the interface and those in the condensate core [10, 19, 21]. In parallel, graph-theoretic analyses have been applied to reveal inhomogeneities in networks of molecular interactions underlying LCD condensates [10– 12, 18, 41]. Building on these observations, we investigate whether molecule-scale conformational properties are systematically related to network features [Fig. 4(a)] within the microstructures of LCD-like condensates. Specifically, we measure the radius of gyration *R*_g_ and shape anisotropy *κ*^2^ of proteins in the condensates. *R*_g_ provides insight into the average molecular size, while *κ*^2^ effectively describes the deviation of polymer shape from a perfect sphere [56–58] [Fig. 4(b)].

**FIG. 4.**
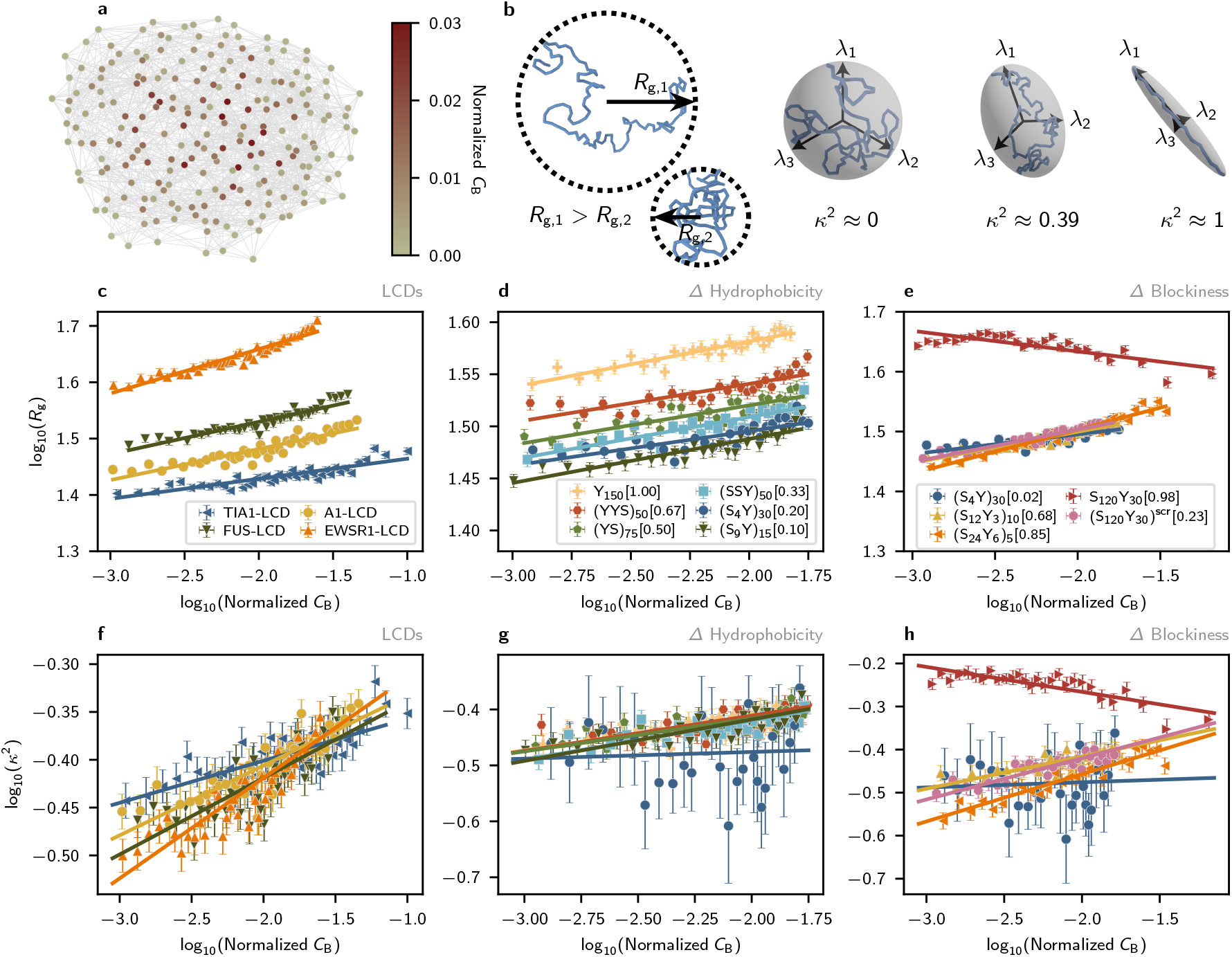
Single-chain radius of gyration *R*_g_ and shape anisotropy *κ*^2^ follow power-law relationships with molecular connectivity in interaction networks. All graph-theoretic analyses are performed by constructing interaction networks using the energy threshold 5*k*_B_*T* to define edges representing long-lived intermolecular associations. **a** A graph of an A1-LCD condensate. Nodes are colored based on their betweenness centrality *C*_B_. **b** (left) Schematic representations of chain radius of gyration *R*_g_. (right) Schematic representations of the eigenvectors *λ* of the gyration tensor and values of the scale-invariant shape anisotropy parameter *κ*^2^ for distinct chain conformations. *κ*^2^ = 0 represents a perfect, radially isotropic sphere, *κ*^2^ ≈ 0.39 corresponds to an ideal-chain conformation, and *κ*^2^ = 1 describes a perfectly anisotropic elongated chain. **c, d, e** Possible power-law relations between the betweenness centrality *C*_B_ and radius of gyration *R*_g_, indicated by linear fits in log_10_–log_10_ space. **f, g, h** Possible power-law relations between the betweenness centrality *C*_B_ and relative shape anisotropy *κ*^2^.

We first compare single-molecule *R*_g_ against molecular betweenness centrality *C*_B_ (normalized; see Methods). Recall that *C*_B_ quantifies the importance of a molecule (i.e., node) based on how often it lies on the shortest paths between other molecules, such that a higher *C*_B_ indicates a more central, globally influential position within the net-work. We find that *R*_g_ versus *C*_B_ in log_10_–log_10_ space yields a positive linear relationship [Fig. 4(c)] for LCDs, suggesting a consistent power-law relationship:

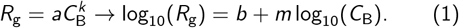

This power-law-like behavior is also observed in condensates formed by uniformly patterned binary YS sequences [Fig. 4(d)]. In both cases, individual macromolecules become more expanded as their network centrality increases, indicating that condensate microstructures reliably encode inhomogeneities at the single-molecule scale. Among LCD sequences, Fig. 4(c) shows that longer chains such as EWSR1 exhibit larger radii of gyration, consistent with expected scaling effects. For uniformly patterned YS sequences of fixed length, *R*_g_ increases monotonically with *C*_B_ with similar linear slopes across all variants [Fig. 4(d)]. Comparing between these sequences reveals that the baseline expansion (intercept) increases with sequence hydrophobicity, resulting in a clear ordering of YS variants by both *R*_g_ and hydrophobic fraction *f*_h_. This trend suggests that the solvent quality of the dense phase, as experienced by individual macromolecules, is not solely determined by intrinsic sequence properties but also by emergent properties of the condensate itself. Indeed, as the solvent quality increases over an order-of-magnitude change in sequence hydrophobicity, chains become more expanded even as their self-associations become more favorable. By opening associative sites to the surrounding environment, this chain expansion promotes the formation of intermolecular associations that appear to strengthen the small-world-like internal architecture [Fig. 2(e)] and the surface tension [Fig. 1(h)].

*R*_g_ profiles for YS sequences with fixed composition and varying patterning appear to corroborate this result [Fig. 4(e)], with all but the blockiest sequence (S_120_Y_30_) having *R*_g_ values collapsing onto the same curve. However, the most blocky sequence S_120_Y_30_ exhibits an almost inverse trend, where increased centrality *C*_B_ is negatively correlated with molecular radii of gyration. This result may be attributed to the packing of hydrophobic poly-Y tails in the hydrophobic core of the micelle formed by S_120_Y_30_.

Similar to *R*_g_, the molecular relative shape anisotropy *κ*^2^ also appears to follow power-law relationships with molecular *C*_B_ for LCDs and non-blocky binary YS sequences, albeit with smaller coefficients of determination *R*^2^ [Fig. 4(f,g)]. *κ*^2^ is a scale-invariant quantity, ranging from *κ*^2^ = 0 at the limit where polymers adopt a spherically isotropic conformation to *κ*^2^ = 1 at the limit where they are completely linear [Fig. 4(b), right panel]. LCD simulations reveal that greater slopes in the *κ*^2^(*C*_B_) relationship correlate with greater small-worldness in the network and greater surface tension [Figs. 4(f), 1(g), and 2(d)], though this trend is not obvious for the YS sequences and likely arises from sequence complexity not captured in the YS variants [Fig. 4(g,h)]. Previous simulation studies have reported the average relative shape anisotropy of individual polyampholytes in condensed phases [19], with dense-phase *κ*^2^ consistently between 0.42 and 0.44 invariant to changes in sequence. However, our results show that LCDs in dense phases exhibit a range of conformations that map onto their connective roles in the condensate microstructure. Compared to the ideal-chain *κ*^2^ = 0.39, molecules with low *C*_B_ adopt slightly collapsed conformations (*κ*^2^ ≈ 0.28) while highly connected hublike molecules become slightly expanded (*κ*^2^ ≈ 0.48).

In summary, we demonstrate the potential to interpret complex single-molecule inhomogeneities in dense phases using principled analyses of condensate microstructures. We predict the presence of continuous quantitative relationships bridging single-molecule conformation and network centrality, whose parameters depend on sequence features and length. Notably, we find that hublike character corresponds to maximal expansion among the conformations assumed by macromolecules in all condensates formed by LCD and non-blocky YS sequences. Simulations of YS variants show that greater sequence hydrophobicities promote chain expansion, stronger small-world-like networking, and greater droplet surface tension. Together, these results show how sequence properties shape the intricate relations between the molecular conformations, mesoscale organization, and macroscopic behaviors of biomolecular condensates.

### Dynamics of molecules in biomolecular condensates are dependent on network topology

In addition to linking molecular conformation and condensate microstructure, we characterize the dynamics of individual molecules inside condensates. In particular, we investigate scaling relationships between the displacement of molecular centers of mass (|*Δ***r**|) and betweenness centrality (*C*_B_). The computation and normalization of these quantities are described in the Methods.

Similar to the conformational properties *R*_g_ and *κ*^2^, we find that normalized molecular displacement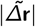 follows an apparent power-law relationship with *C*_B_, exhibiting linear correlations in log_10_–log_10_ space [Fig. 5(a,b,c)]. Given that the LCD sequences have similar hydrophobicities, the LCD data mainly show that molecular size strongly impacts displacement; the shortest sequence TIA1 exhibits the fastest dynamics and the longest sequence EWSR1 exhibits the slowest dynamics [Fig. 5(a)]. YS sequences show that increased sequence hydrophobicity is associated with greater overall molecular displacement when compared at 0.9*T*_c_ [Fig. 5(b)]. This trend suggests that, at a fixed relative distance from the critical temperature, more hydrophobic sequences experience a more favorable environment (i.e., better solvent quality) within the dense phase, promoting increased mobility and shaping the internal dynamics of the condensate. These results reveal that sequence-encoded microstructures are linked to single-molecule dynamics as well as conformational properties, supporting their explanatory power for both static and dynamic condensate phenomena.

**FIG. 5.**
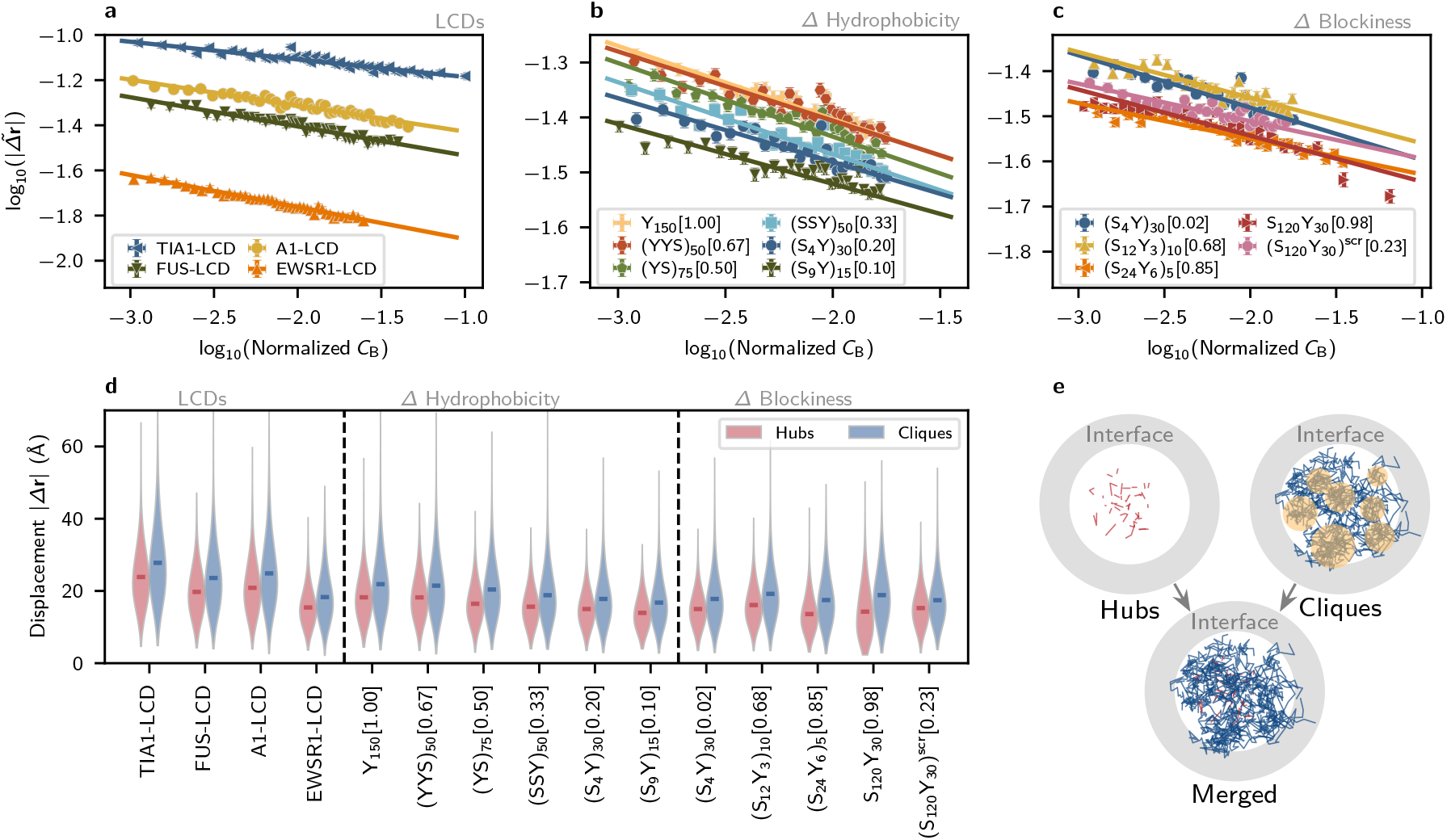
Dynamics of molecules within single-component condensates is correlated with network topological organization. **a,b, c** Normalized instantaneous displacements 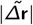 of single chains in condensed phases exhibit strong negative correlations with node betweenness centrality *C*_B_ within molecular graphs. **d** Displacements within consistent time intervals are compared for hub molecules and clique molecules. **f** A 2-dimensional visualization of the trajectories of hub molecules (red) and clique molecules (dark blue) in a FUS-LCD condensate, plotted as lines when their hub or clique-member status is contiguous in time. The phase interface is shown as a gray circular band. Locally confined regions of clique molecules are indicated by yellow circles.

Experimental and simulation studies have characterized the diffusion of LCDs within condensates [59, 60]. More recent experimental studies have shown that the dense phases of FUS–RNA condensates contain “nanodomains”, densely interacting molecular clusters that decrease local diffusivity without a secondary phase separation [40]. To study this effect and its relation to the small-world microstructure, we additionally analyze the local movements of hub and clique molecules in our systems. On short timescales, the displacements of clique molecules are consistently greater than those of hubs in both LCD and YS-variant simulations [Fig. 5(d)]. This can be intuitively explained with the observed degree distribution of hub molecules and clique molecules within interaction networks: hub molecules associate with a greater number of partners, thus confining their motion to a greater extent (Figs. S11–S14). When combined with our previous analysis, which reveals that cliques have longer lifetimes [Fig. 3(d,e)], we conclude that the “faster” motion of clique molecules is best described as a form of local vibration. Indeed, when we trace the displacement of molecules in cliques and hubs, the motion of clique molecules is highly localized (see the 2D projections of molecular motion in Fig. 5(f)). Thus, while cliques experience relatively larger displacements than hubs, these displacements remain confined to microscale regions near the interface. Such confinement of clique motion is reminiscent of the nanodomains described in Ref. 40, as well as of recent experiments reporting highly interactive hydrophobic “hotspots” in A1-LCD condensates [12]. These results support the hypothesis that nanoscale clusters at interfaces might contribute to selective molecular recruitment and confinement through local modulation of material properties.

## DISCUSSION

Macroscopically, biomolecular condensates often appear as homogeneous structures in both experimental and simulated reconstitution. However, evidence suggests that even single-component condensates can exhibit inhomogeneities in their microstructure [10–12, 15, 37]. The molecular networks underlying condensates have also been theorized to play crucial roles in shaping their material properties and functions. Understanding how the material properties of biomolecular condensates arise from complex microstructures and molecular features is thus an emerging area in the field.

In this work, we leverage residue-resolution molecular dynamics simulations alongside graph theory to reveal how single-molecule conformations, condensate microstructure, and emergent material properties are coupled across scales. We systematically characterize the microstructures of condensates formed by the low-complexity domains (LCDs) of key phase-separating proteins (hnRNPA1, FUS, EWSR1, and TIA1) using network-based approaches. To assess the generality of our findings, we also characterize condensate systems composed of binary sequences with varying fractions of hydrophobic Y residues and polar S residues. These YS variant systems represent associative heteropolymers with varying propensities for forming percolated networks within condensates.

Consistent with previous lattice-based simulations of A1LCD [10], we find that condensates formed by biological LCDs and generic non-blocky SY sequences adopt microstructures well described as small-world networks. Notably, we discover that higher sequence hydrophobicities engender more small world-like internal architecture and greater droplet surface tension, directly linking sequence to condensate microstructure and macroscopic material properties. Small-world networks contain two major topological features, “hubs” and “cliques”. Cliques are densely connected groups of nodes (here, molecules) representing fully connected subgraphs within the network. The cliques themselves are efficiently bridged through hub nodes, which are highly connective and reduce average shortest path lengths between arbitrary node pairs in the network. We consistently find that hubs are positioned closer to the condensate center, while the largest cliques are located near the interface. Such spatial organization of network features enables efficient transmission of internal stresses to clique clusters at the interface, potentially driving the increase in effective surface tension. Condensate microstructures, then, can be viewed as spatially embedded networks whose complex properties directly account for emergent material behaviors.

Our work also elucidates how mesoscale inhomogeneities arise from internal network architectures. Hubs and cliques represent two distinct regimes of molecular interactivity within the microstructure: hub molecules are responsible for high global connectivity and are marked by expanded conformations, while clique clusters are nanoscale regions of elevated local associativity marked by interfacial localization and confined molecular motion. In addition to the spatial distinction observed between hubs and cliques, we find that hub molecules and clique molecules have distinct lifetimes: the molecular identities of connective hubs change rapidly, while members of cliques tend to remain in those cliques over longer timescales due to the formation of stable, fully-connected subnetworks. These results are highly counterintuitive. Network hubs are critical for network stability and often form many strong intermolecular associations, but their role is shown to be transient. Network cliques represent molecular clusters at the interface marked by maximal local connectivity, making them sensitive to minor perturbations, yet clique structures are demonstrably more time-stable. Such inverted behavior may be a crucial feature of resilience and function in LCD condensates: transient hubs allow decentralization of the backbone holding the network together, while stable clusters may modulate selective molecular recruitment and dynamic arrest by controlling the material characteristics of the interface. Importantly, we demonstrate that the properties of interfacial clusters can be tuned with sequence design, providing a guiding principle for interpreting condensate aging and engineering condensate function.

At the single-molecule scale, we reconcile reports of conformational heterogeneity and connective heterogeneity in condensates by showing that the conformational properties of macromolecules are highly correlated with their connectivity within network microstructures. Interestingly, the radius of gyration (i.e., average size) and shape anisotropy (i.e., ranging from spherical to linear) of individual molecules are found to follow power-law-like relations to molecular betweenness centrality (i.e., connectivity) within interaction networks. This behavior is conserved across a range of sequence lengths, compositions, and patterns. We find that increasing sequence hydrophobicity does not alter the nature or the slope of the relationship. Instead, increased hydrophobicity leads to greater overall chain expansion, which corresponds to strengthened small-world structure and increased droplet surface tension. Further, we find that sequence hydrophobicity does not appear to alter conformational anisotropy in dense phases but sequence length strongly impacts the degree of chain extension and linearization with increasing connectivity. Lastly, we note that the blockiest sequences (S_24_Y_6_)_5_ and S_120_Y_30_ form micelles instead of phase separating. Curiously, the block-copolymer sequence S_120_Y_30_ exhibits relationships that are inverted relative to all other sequences, and we attribute this behavior to packing and disorder in the hydrophobic micelle core and periphery, respectively. This behavior is not representative of phase-separated macromolecules in solution.

It is striking that molecule-scale physical quantities are strongly correlated with betweenness centrality and not with node degree, the intuitive measure of polymer associativity. Betweenness centrality and degree are naturally weakly correlated (Figs. S7–S10), as a node with a greater number of associations has an increased likelihood of lying along shortest paths, i.e., being globally connective. Indeed, on average, node centrality and degree vary similarly with increasing radial distance from the condensate center (Figs. S15–S18). However, high betweenness centralities do not require high degrees and vice versa. We find that the diverse conformational and dynamic characteristics assumed by single molecules in dense phases are most coherently viewed in a relationship with *C*_B_ and not with degree or radial distance. We take these findings to mean that shortestpath centrality is a critical organizing principle of the condensate microstructure, and further, that inhomogeneities at the single-molecule scale are more strongly related to emergent properties of internal networking than directly to the number of intermolecular bonds or to spatial location within the dense phase. This, along with the observed ubiquity of the small-world topology, corroborates the notion that small-world microstructures give rise to both microscopic properties—through the relations uncovered here— and macroscopic material properties, by enabling efficient internal stress transmission.

In addition to structural properties of macromolecules, we explore their dynamics in the context of the microstructure. We find that molecules become less dynamic as their betweenness centrality increases. This result is intuitive, as more expanded molecules with larger betweenness centralities are subject to greater confinement through dense intermolecular interactions in the condensate environment. Interestingly, this relationship also shows power-law behavior. As expected, we find that smaller molecules exhibit faster motions in the dense phase. Increasing sequence hydrophobicity also leads to greater overall displacements across the range of the relationship, further illuminating how sequence features tune the relation between condensate microstructure and molecular properties.

To our surprise, molecules in interfacial cliques exhibit slightly faster motions than hub molecules. Explicitly tracing their trajectories, however, reveals that clique molecules are spatially confined. This suggests that their movement is primarily characterized by local vibrations, whereas hub molecules exhibit motion across larger regions. Recent work on multicomponent condensates has identified nanodomains within condensates marked by elevated local connectivity, interfacial localization, and reduced molecular diffusion [40], all of which appear to be consistent with the cliques we observe in LCD and YS condensates. Other recent experiments have also reported highly interactive hydrophobic nanoclusters in A1-LCD condensates [12]; although Ref. [12] refers to these regions as “hubs,” their nanoclusters align with the “cliques” found in our systems according to graph-theoretic principles. Alongside these studies, experiments report that pathological liquid-to-gel transitions in condensates originate at their interfaces [31, 61]. Additionally, the formation of interfacial aggregates resembling amyloid fibrils were observed in the early stages of FUS condensate aging [32, 61]. The arresting effects of interfacial aggregation can, in turn, arrest the dynamics of the entire condensate by propagating through the small-world network structure. Thus, the stability of locally constrained, highly interactive molecular clusters near condensate interfaces may be linked to both function and dysfunction, enabling selective molecular recruitment and pathological aggregation.

A key limitation of this work is that the probed condensates are composed solely of disordered protein sequences that engage in transient interactions. In contrast, cellular condensates often include both disordered and structured components that dictate their form and function [62]. The latter can mediate long-lived, high-affinity interactions that shape the underlying interaction networks, as recent studies suggest [63, 64]. Exploring systems with specific binding interactions—such as those involving folded protein domains or RNA—would therefore be an important next step. Nonetheless, our findings show that even simple systems of disordered protein regions exhibit striking microstructural inhomogeneities that give rise to complex biophysical behaviors.

Collectively, our results elucidate the rules by which molecular sequences and features encode the rich internal microstructures of condensates, and, in turn, how microstructures give rise to the complex material behaviors observed of condensates. We demonstrate that LCD-like condensates consistently adopt small-world network microstructures that are directly linked to sequence features such as hydrophobicity and macroscopic properties such as surface tension. Further, we analyze the spatial and temporal characteristics of distinct network features, hubs and cliques, to interpret experimentally observed mesoscale inhomogeneities in the context of the microstructure. We finally uncover quantitative relationships that govern the dis-tribution of molecular conformations and dynamics in dense phases from molecular network connectivity. These findings reveal complex, multiscale structure–property relationships in LCD condensates that provide principles for the rational design of soft materials with complex internal architectures. We anticipate that the framework developed here can be extended to other biomolecular systems to inform the design of synthetic condensates with programmable and interpretable properties.

## METHODS

In this work, we study the single-chain characteristics, interaction network topologies, and surface tensions of single-component condensates formed by prion-like lowcomplexity domains (LCDs) using Mpipi—a residue-level coarse-grained model for disordered proteins [42]. We further study the effect of sequence on multiscale properties by simulating single-component condensates comprising binary sequences of tyrosine (Y) and serine (S) residues. In the binary sequence simulations, we systematically vary sequence hydrophobicity (fraction of Y, *f*_h_) and blockiness (fraction of consecutive residues, *f*_B_) to investigate their impact on the topology of emergent interaction networks, as well as their downstream impact on surface tension as well as molecular conformation and dynamics.

### 1. LCD and Binary sequences

We use the Mpipi model [42] to simulate four biological phase-separating protein sequences: the low-complexity domain of the heterogeneous nuclear ribonucleoprotein hnRNPA1 (A1-LCD), the low-complexity domain of the Fused in Sarcoma protein (FUS-LCD), the low-complexity domain of the RNA-binding protein EWS (EWSR1-LCD), and the low-complexity domain of the T-cell intracellular antigen 1 (TIA1-LCD). The sequences are shown below.

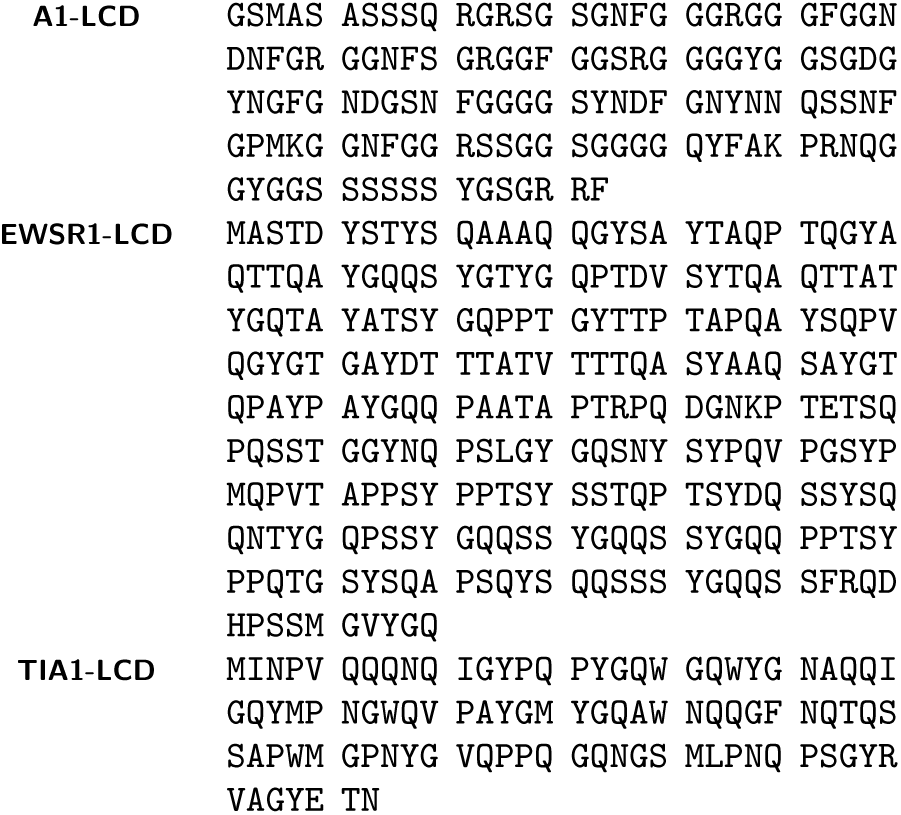
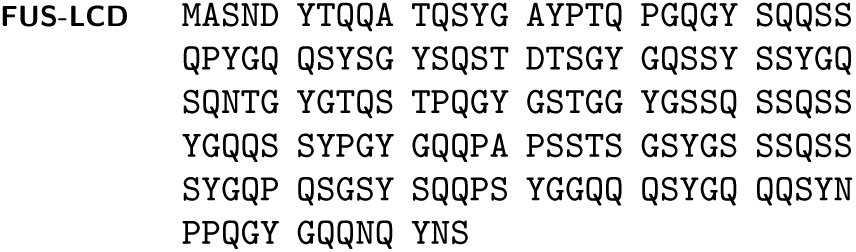

These LCDs are marked by a sequence distribution overrepresented in Glutamine (Q), Serine (S), Glycine (G), and Tyrosine (Y) residues. The polar uncharged residues Q, S, and G act as weakly interactive “spacers” along sequences, serving to segregate highly attractive “sticker” residues (particularly Y) uniformly along the sequence. TIA1-LCD also incorporates tryptophan (W) residues along the sequence that can enable “sticky” interactions with itself and tyrosine (Y) through π–π stacking of aromatic rings.

As for the binary SY sequences, we simulate chains with a constant length *n* = 150 to be close to the length of the LCD sequences described above. Furthermore, 11 sequence variants with different hydrophobic fractions are simulated and analyzed: (S_4_Y)_30_, (S_120_Y_30_)^scr^, (YS)_75_, Y_150_, (S_12_Y_3_)_10_, (S_24_Y_6_)_5_, (YYS)_50_, (S_9_Y)_15_, S_120_Y_30_, (SSY)_50_, and S_150_. Analogous to LCD architectures, each constructed variant distributes hydrophobic Y beads as evenly as possible along the sequence. These sequences represent a range of sequence compositions spanning an order-of-magnitude change in hydrophobicity. Among the sequences, S_150_ has the lowest hydrophobicity *f*_h_ = 0.00 and does not phase separate at *T >* 104 K; we have thus excluded this sequence from the results. Y_150_ has the largest hydrophobicity *f*_h_ = 1.00, and correspondingly, the highest critical temperature *T*_c_ = 945 K. In between, we cover a wide range of hydrophobicities: *f*_h_ = 0.10 for (S_9_Y)_15_; *f*_h_ = 0.20 for (S_4_Y)_30_, (S_4_Y)^scr^, (S_12_Y_3_)_10_, (S_24_Y_6_)_5_, and S_120_Y_30_; *f*_h_ ≈ 0.33 for (SSY)_50_; *f*_h_ ≈ 0.67 for (YYS)_50_. Since *f*_h_ = 0.20 is similar to the hydrophobicity of LCD sequences (*f*_h_ ≈ 0.14), we vary sequence blockiness at this fixed composition, and we also scramble a sequence to further examine the effects of sequence patterning. This scrambled sequence is denoted (S_120_Y_30_)^scr^.

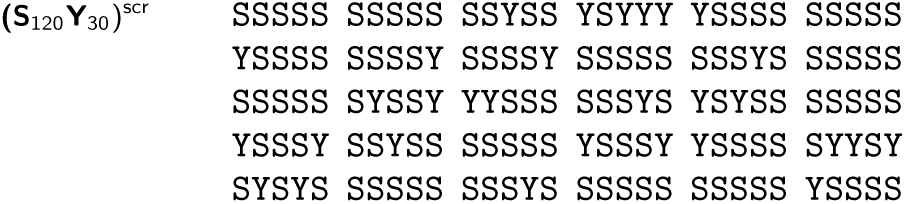

The blockiness *f*_B_ is quantified by calculating the ratio of the number of actual Y–S and S–Y bonds *B*_act_ over the possible maximum number of Y–S and S–Y bonds *B*_max_,

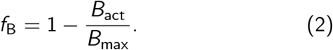

When *f*_B_ → 1, it means that there are fewer Y–S and S–Y bonds, indicating the increased blockiness. On the opposite side, the least blocky sequence maximizes the number of Y–S and S–Y bonds, thus *f*_B_ → 0.

### 2. Mpipi model

In Mpipi, each protein residue is represented by a single interaction site/bead. Each bead has an assigned mass, charge, molecular diameter, and other interaction parameters. The potential energy in the Mpipi model is taken as the sum of bonded and non-bonded interaction terms.

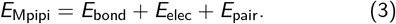

Specifically, beads are bonded via harmonic springs.

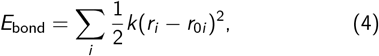

where the bond spring constant *k* is 19.2 kcal mol^−1^Å^2^ and the equilibrium bond length is 3.81Å.

Non-bonded interactions encompass long-ranged electrostatics, which are captured via a Coulomb term with Debye– Hückel screening,

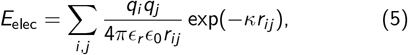

where *ϵ*_r_ = 80 is the relative dielectric constant of water and *ϵ*_0_ is the electric constant. The Debye screening length is set to *κ*^−1^ = 7.95 Å with a Coulomb cutoff of 35 Å [42].

Non-bonded interactions also include short-ranged pairwise contacts, which are modeled via the Wang–Frenkel potential [65],

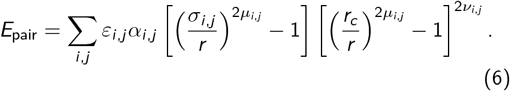

All model parameters are discussed in detail in Refs. 42 and 38 and are provided in our GitHub repository (see Data Availability Statement). In Mpipi, the solvent is modeled implicitly, and the model was parameterized by combining bioinformatics data and atomistic potentials-of-mean force calculations. Previous work has demonstrated that Mpipi accurately captures both single-chain properties and collective phase behaviors of disordered proteins [38, 42, 66].

### 3. Molecular dynamics simulations

Implicit-solvent simulations are conducted in the *NV T* ensemble using the LAMMPS package [67].

First, *NPT* simulations are performed to accelerate the condensate formation process during the initial steps. A Berendsen barostat is used to apply an isotropic external pressure to the particles in each simulation cell, effectively overcoming the nucleation barrier and compressing the polymer chains into a condensed state. LCD simulations are compressed with a fixed isotropic pressure set to 100 atmospheres for 120,000 timesteps (*dt* = 10 fs) with a pressure damping parameter of 10,000 *dt*. YS sequence variants are subject to a time-varying pressure that increased from 50 atmospheres to 100 atmospheres over a period of 30,000 timesteps (*dt* = 10fs) with a pressure damping parameter of 100,000 *dt*.

Production runs are then performed at 0.9*T*_c_ and 0.95*T*_c_. The integration timestep is set to *dt* = 10 fs, and systems are simulated for 1 *µ*s after condensate formation for equilibrium sampling. 1000 frames are sampled uniformly along equilibrium trajectories for each sequence. For each sampled frame in both simulation types, dense-phase centers of mass and single-molecule conformations are obtained using OVITO [68].

### 4. Construction of interaction graphs

Interaction matrices representative of single static frames are constructed from particle position data, and we use an energetic criterion to ensure that the interaction energy of two chains *a* and *b* exceeds the thermal energy, i.e., *E*_pair,*ab*_ < − 5*k*_B_*T*, when recording an interaction. For *N* condensed polymers, a 2-dimensional interaction matrix (adjacency matrix) *M* = *N* × *N* is constructed, where *M*_*ab*_ is assigned as follows:

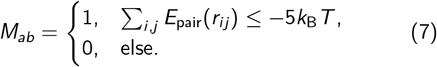

Here *i* and *j* index over each bead (residue/monomer) along respective protein chains *a* and *b*, and *r*_*ij*_ is the distance between monomers *a* and *b*. Finally, graph structures are generated with the NetworkX python package [69] using binary interaction matrices as adjacency matrices *M*. Each node in a frame’s representative graph represents a single protein chain, and unweighted, undirected edges are drawn between nodes if an attractive interaction between them is observed in that frame according to the above criteria.

Interaction networks are studied for small-world-like topologies by finding node betweenness centralities *C*_B_ and calculating the small-world coefficients *σ*_sw_ and *ω*_sw_ [49, 51, 53]. The betweenness centrality *C*_B_ of a node *i* is found and normalized via the NetworkX betweenness_centrality() utility and is computed as follows:

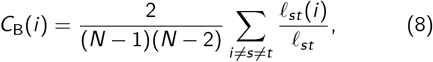

where the pair (*s, t*) enumerates over all node pairs in the graph (excluding *i*), *ℓ*_*st*_ is the total number of shortest paths between *s* and *t*, and *ℓ*_*st*_(*i*) is the number of shortest paths that flow through node *i*. The normalization coefficient is the inverse of the binomial coefficient 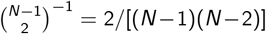 for a graph with *N* − 1 nodes, enumerating over all combinations of node pairs excluding *i*. Betweenness centralities are normalized to facilitate comparison between graphs of systems of differing sizes, as the summation suggests that it is a metric that scales with the number of nodes *N*.

Graphs are generally considered to have small-world topologies if neighbors of any given node are highly connected to each other, if shortest pathlengths between any given pair of nodes are low, and if the graph is sparse [49]. Both *σ*_sw_ and *ω*_sw_ serve as estimators of the “smallworldness” of a given graph by comparing its average shortest pathlength *L* = ⟨*ℓ*_min_⟩ and its average clustering coefficient *C* to the same quantities *C*_rand_ and *L*_rand_ found for a series of Erdös–Rényi random graphs, and *C*_latt_ and *L*_latt_ for equivalent lattice graphs :

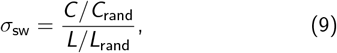

and

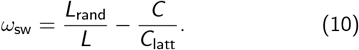

It is commonly recognized that “small-worldness” in a graph corresponds to *σ*_sw_ *>* 1 and *ω*_sw_ ≈ 0 [51–53]. Small-world network topologies are marked by high clustering and low average pathlengths, i.e., that individual “subcommunities” of nodes exist within the graph that are closely connected to each other and that particular nodes serve as highly-connected hubs bridging each subcommunity together in an efficient manner. The flow of information or impulses within these small-world networks are thus efficient with minimal loss in fidelity.

The betweenness centrality *C*_B_ of a node *i* serves as a metric on its hub-like connectivity, measuring the number of shortest paths between any arbitrary node pair that flows through *i*. For each sampled frame, the ten graph nodes with the highest normalized *C*_B_ are selected as “hubs,” and the ten largest cliques are selected as the “subcommunities” of closely-connected nodes. A maximal clique for any node *i* is defined as the largest fully-connected subgraph containing *i* within the graph of interest; maximal cliques are found via the NetworkX *find_cliques()* utility and only reported if fewer than three nodes within the clique are members of an existing reported clique. In simulations where fewer than ten cliques existed, as is possible in the 125-chain HP model simulations, all independent cliques larger than a triangle (3 nodes) are reported.

### 5. Spatial organization within simulated condensates

To study the spatial distribution of topological features from condensate simulations, continuous trajectory samples comprising 20% of total production runs (LCD: 200 ns of 1 *µ*s; HP sequences: 2 × 10^7^ *dτ* of 10^8^ *dτ*) are used, and molecular hub and clique statuses are recorded for each frame. Average radial mass density profiles are generated for each simulation to obtain phase interfaces in tandem with data on the radial distribution of hubs and cliques.

In each sampled frame, all particle masses and radial distances from the dense-phase center of mass are collected and aggregated, and the mass densities are computed by radial binning. Data on the spatial localization of hubs and cliques are recorded by locating hub and clique molecules within the condensate, computing their centers of mass, determining the radial distances between molecular centers of mass and dense-phase centers of mass, and binning. Sigmoid functions are used to fit each radial mass density profile with the *scipy.optimize.curve_fit()* utility [70] to quantitatively define interfacial boundaries. The radial bounds of the interface correspond to the radial distances where the mass density is 95% and 5% of the stable dense-phase sigmoid fit value, capturing most of the region of change. Finally, distances in radial distributions are normalized by their corresponding simulation’s upper (dilute-phase) interfacial boundary in order to facilitate comparison between systems.

### 6. Graph dynamics of the simulated condensate

To understand the time variance of topological organizations, we compute the timescales associated with the presence of hubs and cliques. The same continuous trajectory samples described in the previous section 5 are used. In each frame, the molecular indices corresponding to the ten nodes with the highest betweenness centralities and the molecular indices corresponding to the members of the 10 largest cliques are recorded. The frequency of single-molecule hub or clique status is computed as the number of frames where individual molecules are labeled as hubs or as associated with cliques, respectively. These frequencies are then normalized by the total number of sampled frames in each continuous trajectory sample.

### 7. Conformational analysis

To analyze the structural and conformational properties of single polymers in our simulations, we computed singlemolecule radii of gyration *R*_g_ and relative shape anisotropies *κ*^2^. These metrics are further employed to quantify changes in IDP conformational properties with respect to molecular connectivities within constructed interaction networks. Entire 1 *µ*s or 10^8^ *dτ* trajectories are sampled at intervals of 20 ns or 2 × 10^6^ *dτ* (i.e., every 20th frame). At each sampled frame, graph analyses are performed as described above to compute betweenness centralities *C*_B_ for each molecule. OVITO is used to obtain *R*_g_ values and diagonalized gyration tensors *S* for each individual molecule:

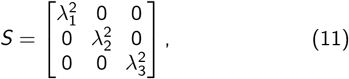

where eigenvalues *λ*_1_, *λ*_2_, *λ*_3_ are the principal components of the molecular gyration tensor. Relative shape anisotropies *κ*^2^ are then obtained via

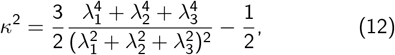

which is bounded between 0 and 1.

### 8. Molecular motion through single-molecule displacement

Heterogeneities in molecular movements within condensed phases are studied by measuring single-molecule displacements at 1 ns (10^5^ *dτ* for HP sequences) intervals, the minimum timestep between static frames in our trajectories. As in our conformational analyses, we sample frames across trajectories of length 1 *µ*s (or 10^8^ *dτ*) at intervals of 20 ns (or 2 ×10^6^ *dτ*). For each of these frames, sampled at some timestep *t*, we compute the center of mass **r**_*i*,COM_(*t*) of each molecule *i*. The magnitudes of “instantaneous” displacements |*Δ***r**_*i*_|are obtained by averaging the differentials of the **r**_*i*,COM_ from frames 1 ns (or 10^5^ *dτ*) before and after the sampled frame at *t*, i.e., by computing **r**_*i*,COM_(*t* − 1) and **r**_*i*,COM_(*t* + 1), respectively:

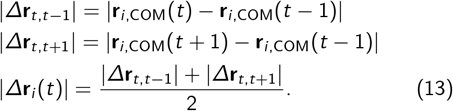

Molecular displacements|*Δ***r**| are also normalized by the length of the corresponding polymer’s linear chain conformation to obtain 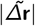 for comparison. Graph analyses are then performed to relate single-molecule displacement to molecular connectivity and topological status.

### 9. Surface tension calculation

Surface tension between two phases of different densities is usually described by the Kirkwood–Buff formalism [71, 72], considering that the presence of an interface provides anisotropy to the overall pressure tensor. Thus, following the definition of mechanical equilibrium, the surface tension is given by:

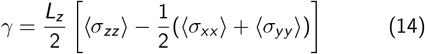

where, *L*_*z*_ is the length of the slab, and *σ*_*ii*_ are the ensemble averages of the diagonal components of the pressure tensor.

However, the formalism is built upon the assumption of a sharp interface, which breaks down as a condensate’s critical temperature is approached. At that limit, the interface become more diffuse and harder to define, yielding incorrect surface tension estimations through the Kirkwood-Buff formalism. To overcome this difficulty, we employ a stress-profile method calculated from per-atom stresses using the virial force contribution equation [73]. This derivation is possible because anisotropy of the diagonal components of the pressure tensors are found at the dense-dilute interface:

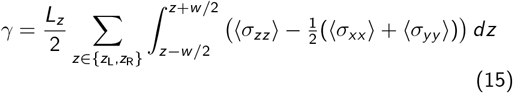

Here *z*_L_, *z*_R_ correspond to the left and right interfaces of the dense phase in the slab geometry, with an equal thickness *w*.

## ACKNOWLEDGMENTS

The simulations reported on in this work were substantially performed using the Princeton Research Computing resources at Princeton University, which is a consortium of groups led by the Princeton Institute for Computational Science and Engineering (PICSciE) and Office of Information Technology. The authors thank Nathaniel Hess for invaluable feedback on the manuscript. The authors also thank other members of the Joseph Group for valuable discussions during various stages of manuscript preparation. D.T. acknowledges research support from the Hewlett Foundation and the New Jersey Alliance for Clinical and Translational Science (NJ ACTS), coordinated through Princeton’s Office of Undergraduate Research. J.A.J. acknowledges research support from the Chan Zuckerberg Initiative DAF (an advised fund of Silicon Valley Community Foundation; grant 2023-332391), and the National Institute Of General Medical Sciences of the National Institutes of Health under Award Number R35GM155259. The content is solely the responsibility of the authors and does not necessarily represent the official views of the National Institutes of Health and other sponsors. This research was partially supported by the National Science Foundation (NSF) through the Princeton University (PCCM) Materials Research Science and Engineering Center DMR-2011750 and in part by grant NSF PHY-2309135 and the Gordon and Betty Moore Foundation Grant No. 2919.02 to the Kavli Institute for Theoretical Physics (KITP).

## AUTHOR CONTRIBUTIONS

D.T., D.A. and J.A.J. conceived the study. D.T., D.A. and P.L.G. performed simulations and analyses. D.T. and D.A. curated and visualized data. D.T., D.A., P.L.G. and J.A.J. wrote and edited the manuscript. D.T. and J.A.J. acquired research funding. D.A. and J.A.J. supervised the research.

## DATA AVAILABILITY STATEMENT

The data supporting the findings in this study, as well as sample simulation input and output files, are available at the Joseph Group GitHub repository: https://github.com/josephresearch/LCD_Network.

## References

[1] C. P. Brangwynne, C. R. Eckmann, D. S. Courson, A. Rybarska, C. Hoege, J. Gharakhani, F. Jülicher, and A. A. Hyman, Germline p granules are liquid droplets that localize by controlled dissolution/condensation, Science 324, 1729 (2009).

[2] A. A. Hyman, C. A. Weber, and F. Jülicher, Liquid-liquid phase separation in biology, Annu. Rev. Cell Dev. Biol. 30, 39 (2014).

[3] Y. Shin and C. P. Brangwynne, Liquid phase condensation in cell physiology and disease, Science 357, eaaf4382 (2017).

[4] D. Zwicker, O. W. Paulin, and C. ter Burg, Physics of droplet regulation in biological cells, arXiv preprint 2501.13639 (2025).

[5] P. Li, S. Banjade, H. Cheng, S. Kim, B. Chen, L. Guo, M. Llaguno, J. V. Hollingsworth, D. S. King, S. F. Banani, P. S. Russo, Q. Jiang, B. T. Nixon, and M. K. Rosen, Phase transitions in the assembly of multivalent signalling proteins, Nature 483, 336 (2012).

[6] R. V. Pappu, S. R. Cohen, F. Dar, M. Farag, and M. Kar, Phase transitions of associative biomacromolecules, Chem. Rev. 123, 8945 (2023).

[7] J. P. Brady, P. J. Farber, A. Sekhar, Y.-H. Lin, R. Huang, A. Bah, T. J. Nott, H. S. Chan, A. J. Baldwin, J. D. Forman-Kay, and L. E. Kay, Structural and hydrodynamic properties of an intrinsically disordered region of a germ cell-specific protein on phase separation, PNAS 114, E8194 (2017).

[8] J. R. Espinosa, J. A. Joseph, I. Sanchez-Burgos, A. Garaizar, D. Frenkel, and R. Collepardo-Guevara, Liquid network connectivity regulates the stability and composition of biomolecular condensates with many components, PNAS 117, 13238 (2020).

[9] T. Mittag and R. V. Pappu, A conceptual framework for understanding phase separation and addressing open questions and challenges, Mol. Cell 82, 2201 (2022).

[10] M. Farag, S. R. Cohen, W. M. Borcherds, A. Bremer, T. Mittag, and R. V. Pappu, Condensates formed by prionlike low-complexity domains have small-world network structures and interfaces defined by expanded conformations, Nat. Commun. 13, 7722 (2022).

[11] F. Dar, S. R. Cohen, D. M. Mitrea, A. H. Phillips, G. Nagy, W. C. Leite, C. B. Stanley, J.-M. Choi, R. W. Kriwacki, and R. V. Pappu, Biomolecular condensates form spatially inhomogeneous network fluids, Nat. Commun. 15, 3413 (2024).

[12] T. Wu, M. R. King, Y. Qiu, M. Farag, R. V. Pappu, and M. D. Lew, Single-fluorogen imaging reveals distinct environmental and structural features of biomolecular condensates, Nat. Phys. (2025).

[13] A. Bremer, M. Farag, W. M. Borcherds, I. Peran, E. W. Martin, R. V. Pappu, and T. Mittag, Deciphering how naturally occurring sequence features impact the phase behaviours of disordered prion-like domains, Nat. Chem. 14, 196 (2022).

[14] F. W. Wiegel, A network model for viscoelastic fluids, Physica 42, 156 (1969).

[15] A. N. Semenov and M. Rubinstein, Thermoreversible gelation in solutions of associative polymers. 2. linear dynamics, Macromolecules 31, 1386 (1998).

[16] D. J. Bauer, L. S. Stelzl, and A. Nikoubashman, Singlechain and condensed-state behavior of hnrnpa1 from molecular simulations, J. Chem. Phys. 157, 154903 (2022).

[17] J. Wang, D. S. Devarajan, A. Nikoubashman, and J. Mittal, Conformational properties of polymers at droplet interfaces as model systems for disordered proteins, ACS Macro Lett. 12, 1472 (2023).

[18] J. C. Shillcock, C. Lagisquet, J. Alexandre, L. Vuillon, and J. H. Ipsen, Model biomolecular condensates have heterogeneous structure quantitatively dependent on the interaction profile of their constituent macromolecules, Soft Matter 18, 6674 (2022).

[19] J. Wang, D. S. Devarajan, K. Muthukumar, Y. C. Kim, A. Nikoubashman, and J. Mittal, Sequence-dependent conformational transitions of disordered proteins during condensation, Chem. Sci. 15, 20056 (2024).

[20] D. S. Devarajan, J. Wang, B. Szała-Mendyk, S. Rekhi, A. Nikoubashman, Y. C. Kim, and J. Mittal, Sequence-dependent material properties of biomolecular condensates and their relation to dilute phase conformations, Nat. Commun. 15, 1912 (2024).

[21] D. J. Bauer and A. Nikoubashman, The conformations of protein chains at the interface of biomolecular condensates, Nat. Commun. 15, 9975 (2024).

[22] L. K. Davis, A. J. Baldwin, and P. Pearce, Mesoscopic heterogeneity in biomolecular condensates from sequence patterning, arXiv preprint 2502.14587 (2025).

[23] M. Feric, N. Vaidya, T. S. Harmon, D. M. Mitrea, L. Zhu, T. M. Richardson, R. W. Kriwacki, R. V. Pappu, and C. P. Brangwynne, Coexisting liquid phases underlie nucleolar sub-compartments, Cell 165, 1686 (2016).

[24] L.-P. Bergeron-Sandoval, S. Kumar, H. K. Heris, C. L. A. Chang, C. E. Cornell, S. L. Keller, P. François, A. G. Hendricks, A. J. Ehrlicher, R. V. Pappu, and S. W. Michnick, Endocytic proteins with prion-like domains form viscoelastic condensates that enable membrane remodeling, PNAS 118, e2113789118 (2021).

[25] L. M. Jawerth, M. Ijavi, M. Ruer, S. Saha, M. Jahnel, A. A. Hyman, F. Jülicher, and E. Fischer-Friedrich, Salt-dependent rheology and surface tension of protein condensates using optical traps, Phys. Rev. Lett. 121, 258101 (2018).

[26] I. Alshareedah, M. M. Moosa, M. Pham, D. A. Potoyan, and P. R. Banerjee, Programmable viscoelasticity in proteinrna condensates with disordered sticker-spacer polypeptides, Nat. Commun. 12, 6620 (2021).

[27] D. Michieletto and M. Marenda, Rheology and viscoelasticity of proteins and nucleic acids condensates, JACS Au 2, 1506 (2022).

[28] I. Alshareedah, W. M. Borcherds, S. R. Cohen, A. Singh, A. E. Posey, M. Farag, A. Bremer, G. W. Strout, D. T. Tomares, R. V. Pappu, T. Mittag, and P. R. Banerjee, Sequence-specific interactions determine viscoelasticity and ageing dynamics of protein condensates, Nat. Phys. 20, 1482 (2024).

[29] D. Sundaravadivelu Devarajan and J. Mittal, Sequence-encoded spatiotemporal dependence of viscoelasticity of protein condensates using computational microrheology, JACS Au 4, 4394 (2024).

[30] D. Aierken and J. A. Joseph, Accelerated simulations reveal physicochemical factors governing stability and composition of rna clusters, J. Chem. Theory Comput. 20, 10209 (2024).

[31] Y. Shen, A. Chen, W. Wang, Y. Shen, F. S. Ruggeri, S. Aime, Z. Wang, S. Qamar, J. R. Espinosa, A. Garaizar, P. St George-Hyslop, R. Collepardo-Guevara, D. A. Weitz, D. Vigolo, and T. P. J. Knowles, The liquid-to-solid transition of fus is promoted by the condensate surface, PNAS 120, e2301366120 (2023).

[32] C. He, C. Y. Wu, W. Li, and K. Xu, Multidimensional superresolution microscopy unveils nanoscale surface aggregates in the aging (milano) of fus condensates, J. Am. Chem. Soc. 145, 24240 (2023).

[33] A. Garaizar, J. R. Espinosa, J. A. Joseph, G. Krainer, Y. Shen, T. P. J. Knowles, and R. Collepardo-Guevara, Aging can transform single-component protein condensates into multiphase architectures, PNAS 119, e2119800119 (2022).

[34] W. Borcherds, A. Bremer, M. B. Borgia, and T. Mittag, How do intrinsically disordered protein regions encode a driving force for liquid–liquid phase separation?, Curr. Opin. Struct. Biol. 67, 41 (2021).

[35] J. Wang, J. M. Choi, A. S. Holehouse, H. O. Lee, X. Zhang,M. Jahnel, S. Maharana, R. Lemaitre, A. Pozniakovsky, D. Drechsel, I. Poser, R. V. Pappu, S. Alberti, and A. A. Hyman, A molecular grammar underlying the driving forces for phase separation of prion-like rna-binding proteins, Cell 174, 688 (2018).

[36] E. W. Martin, A. S. Holehouse, I. Peran, M. Farag, J. J. Incicco, A. Bremer, C. R. Grace, A. Soranno, R. V. Pappu, and T. Mittag, Valence and patterning of aromatic residues determine the phase behavior of prion-like domains, Science 367, 694 (2020).

[37] A. N. Semenov and M. Rubinstein, Thermoreversible gelation in solutions of associative polymers. 1. statics, Macromolecules 31, 1373 (1998).

[38] M. J. Maristany, A. A. Gonzalez, J. R. Espinosa, J. Huertas, R. Collepardo-Guevara, and J. A. Joseph, Decoding phase separation of prion-like domains through data-driven scaling laws, eLife 13, RP99068 (2025).

[39] S. Rekhi, D. S. Devarajan, M. P. Howard, Y. C. Kim, A. Nikoubashman, and J. Mittal, Role of strong localized vs weak distributed interactions in disordered protein phase separation, The Journal of Physical Chemistry B 127, 3829 (2023).

[40] G. Gao, E. R. Sumrall, and N. G. Walter, Single molecule tracking reveals nanodomains in biomolecular condensates, bioRxiv (2024).

[41] Z.-S. Yan, Y.-Q. Ma, and H.-M. Ding, Unveiling the multicomponent phase separation through molecular dynamics simulation and graph theory, J. Chem. Phys. 160 (2024).

[42] J. A. Joseph, A. Reinhardt, A. Aguirre, P. Y. Chew, K. O. Russell, J. R. Espinosa, A. Garaizar, and R. Collepardo-Guevara, Physics-driven coarse-grained model for biomolecular phase separation with near-quantitative accuracy, Nat. Comp. Sci. 1, 732 (2021).

[43] A. Statt, H. Casademunt, C. P. Brangwynne, and Z. Panagiotopoulos, Model for disordered proteins with strongly sequence-dependent liquid phase behavior, J. Chem. Phys. 152 (2020).

[44] G. A. Chapela, G. Saville, S. M. Thompson, and J. S. Rowlinson, Computer simulation of a gas–liquid surface. part 1, J. Chem. Soc., Faraday Trans. 2 73, 1133 (1977).

[45] J. S. Rowlinson and B. Widom, Molecular theory of capillarity (Dover, Mineola, New York, 2013).

[46] C. N. Johnson, K. A. Sojitra, E. J. Sohn, A. K. Moreno-Romero, A. Baudin, X. Xu, J. Mittal, and D. S. Libich, Insights into mol. diversity within the fus/ews/taf15 protein family: Unraveling phase separation of the n-terminal low-complexity domain from rna-binding protein ews, J. Am. Chem. Soc. 146, 8071 (2024).

[47] Y. Lin, S. L. Currie, and M. K. Rosen, Intrinsically disordered sequences enable modulation of protein phase separation through distributed tyrosine motifs, J. Biol. Chem 292, 19110 (2017).

[48] G. L. Dignon, W. Zheng, Y. C. Kim, R. B. Best, and J. Mittal, Sequence determinants of protein phase behavior from a coarse-grained model, PLoS Comput. Biol. 14, e1005941 (2018).

[49] D. Watts and S. Strogatz, Collective Dynamics of Small-World Networks, Nature 393, 440 (1998).

[50] D. J. Watts, Small Worlds (Princeton University Press, Princeton, NJ, 1999).

[51] M. D. Humphries, K. Gurney, and T. J. Prescott, The brainstem reticular formation is a small-world, not scale-free, network, Proc. R. Soc. B 273, 503 (2006).

[52] M. D. Humphries and K. Gurney, Network ‘small-worldness’: A quantitative method for determining canonical network equivalence, PLoS One 3, e0002051 (2008).

[53] Q. K. Telesford, K. E. Joyce, S. Hayasaka, J. H. Burdette, and P. J. Laurienti, The ubiquity of small-world networks, Brain Connect. 1, 367 (2001).

[54] O. Sporns and J. D. Zwi, The small world of the cerebral cortex, Neuroinformatics 2, 145 (2004).

[55] M. Newman, Networks (Oxford University Press, 2018).

[56] D. N. Theodorou and U. W. Suter, Shape of unperturbed linear polymers: polypropylene, Macromolecules 18, 1206 (1985).

[57] H. ArkIn and W. Janke, Gyration tensor based analysis of the shapes of polymer chains in an attractive spherical cage, J. Chem. Phys. 138 (2013).

[58] D. Aierken and M. Bachmann, Impact of bending stiffness on ground-state conformations for semiflexible polymers, J. Chem. Phys. 158 (2023).

[59] K. A. Burke, A. M. Janke, C. L. Rhine, and N. L. Fawzi, Residue-by-residue view of in vitro fus granules that bind the c-terminal domain of rna polymerase ii, Mol. Cell 60, 231 (2015).

[60] W. Zheng, G. L. Dignon, N. Jovic, X. Xu, R. M. Regy, N. L. Fawzi, Y. C. Kim, R. B. Best, and J. Mittal, Molecular details of protein condensates probed by microsecond long atomistic simulations, J. Phys. Chem. B 124, 11671 (2020).

[61] L. Emmanouilidis, E. Bartalucci,, Y. Kan, M. Ijavi, M. E. Pérez, P. Afanasyev, D. Boehringer, J. Zehnder, S. H. Parekh, M. Bonn, T. C. T. Michaels, T. Wiegand, and F. H.-T. Allain, A solid beta-sheet structure is formed at the surface of fus droplets during aging, Nat. Chem. Bio. 20, 1044 (2024).

[62] N. Hess and J. A. Joseph, Structured protein domains enter the spotlight: modulators of biomolecular condensate form and function, Trends Biochem. Sci. (2025).

[63] C. Dollinger, E. Potolitsyna, A. G. Martin, A. Anand, G. K. Datar, J. D. Schmit, and J. A. Riback, Nanometer condensate organization in live cells derived from partitioning measurements, bioRxiv (2025).

[64] D. Aierken, V. Zhang, R. Sealfon, J. C. Marecki, K. D. Raney, A. S. Gladfelter, J. A. Joseph, and C. A. Roden, Biomolecular condensates control and are defined by rna-rna interactions that arise in viral replication, bioRxiv (2024).

[65] X. Wang, S. Ramírez-Hinestrosa, J. Dobnikar, and D. Frenkel, The lennard-jones potential: when (not) to use it, Phys. Chem. Chem. Phys. 22, 10624 (2020).

[66] J. M. Lotthammer, G. M. Ginell, D. Griffith, R. Emenecker, and A. S. Holehouse, Direct prediction of intrinsically disordered protein conformational properties from sequence, Nat. Methods 21, 456 (2024).

[67] A. P. Thompson, H. M. Aktulga, R. Berger, D. S. Bolintineanu, W. M. Brown, P. S. Crozier, P. J. in ‘t Veld, A. Kohlmeyer, S. G. Moore, T. D. Nguyen, R. Shan, M. J. Stevens, J. Tranchida, C. Trott, and S. J. Plimpton, LAMMPS - a flexible simulation tool for particle-based materials modeling at the atomic, meso, and continuum scales, Comp. Phys. Comm. 271, 108171 (2022).

[68] A. Stukowski, Visualization and analysis of atomistic simulation data with OVITO-the Open Visualization Tool, Model. Simul. Mater. Sci. Eng. 18 (2010).

[69] A. A. Hagberg, D. A. Schult, and P. J. Swart, Exploring network structure, dynamics, and function using networkx, in Proceedings of the 7th Python in Science Conference, edited by G. Varoquaux, T. Vaught, and J. Millman (Pasadena, CA USA, 2008) pp. 11 – 15.

[70] P. Virtanen, R. Gommers, T. E. Oliphant, M. Haberland, T. Reddy, D. Cournapeau, E. Burovski, P. Peterson, W. Weckesser, J. Bright, S. J. van der Walt, M. Brett, J. Wilson, K. J. Millman, N. Mayorov, A. R. J. Nelson, E. Jones, R. Kern, E. Larson, C. J. Carey, I. Polat, Y. Feng, E. W. Moore, J. VanderPlas, D. Laxalde, J. Perktold, R. Cimrman, I. Henriksen, E. A. Quintero, C. R. Harris, A. M. Archibald, A. H. Ribeiro, F. Pedregosa, P. van Mulbregt, and SciPy 1.0 Contributors, SciPy 1.0: Fundamental Algorithms for Scientific Computing in Python, Nat. Methods 17, 261 (2020).

[71] K. Silmore, M. Howard, and A. Panagiotopoulos, Vapour– liquid phase equilibrium and surface tension of fully flexible lennard–jones chains, Mol. Phys. (2017).

[72] D. T. Walton, J. Rowlinson, and J. Henderson, The pressure tensor at the planar surface of a liquid, Mol. Phys. (1983).

[73] A. P. Thompson, S. J. Plimpton, and W. Mattson, General formulation of pressure and stress tensor for arbitrary many-body interaction potentials under periodic boundary conditions, J. Chem. Phys. (2009).

