## Supporting Information for "Biomolecular condensate microstructure couples molecular and mesoscale properties"

### Supporting information for:
### Biomolecular condensate microstructure couples molecular and mesoscale properties

Daniel Tan,[1,*] Dilimulati Aierken,[1,2,*] Pablo L. Garcia,[1] and Jerelle A. Joseph[1,2,†]

[1]*Department of Chemical and Biological Engineering, Princeton University, Princeton, NJ 08544, USA*
[2]*Omenn–Darling Bioengineering Institute, Princeton University, Princeton, NJ 08544, USA*

(Dated: July 19, 2025)

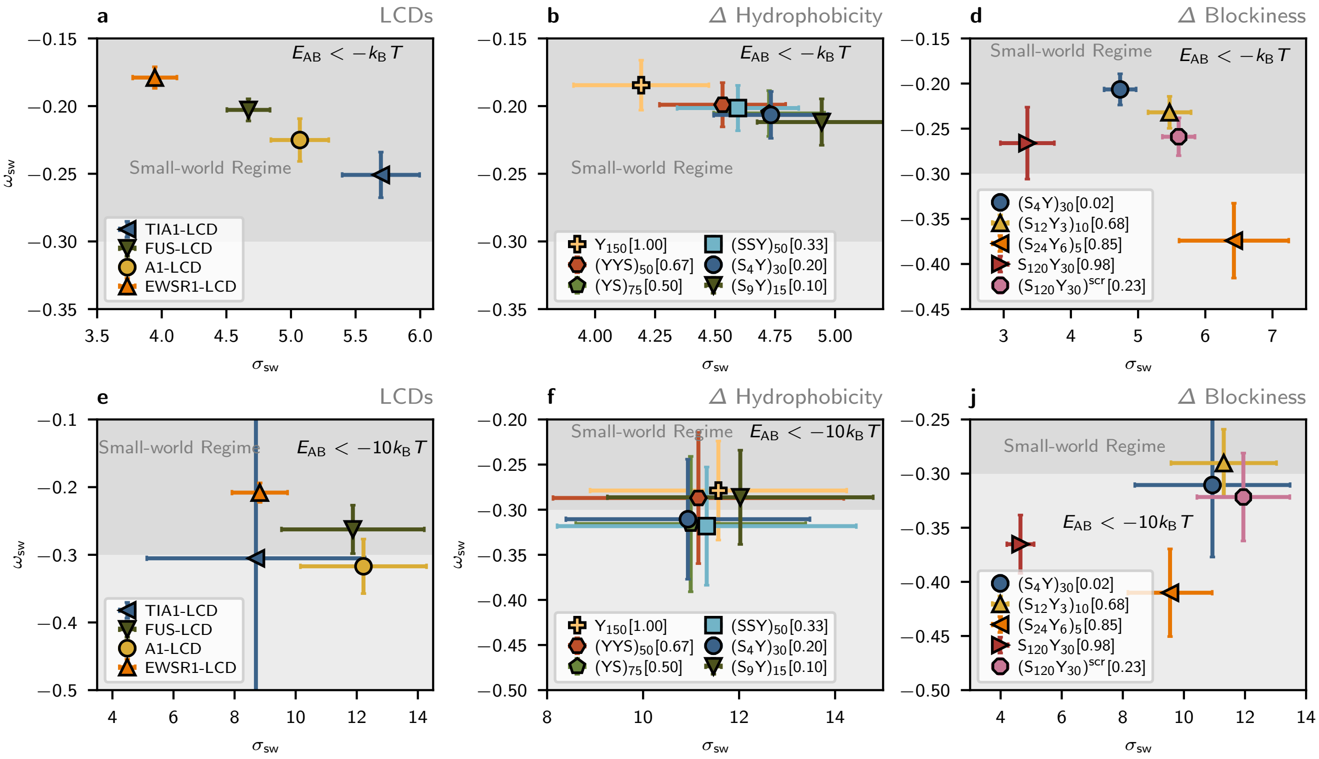


FIG. S1. **a,e** Small-world parameters $\sigma_{sw}$ and $\omega_{sw}$ for single-component LCD condensates with energy threshold $E_{AB} < -k_B T$ and $E_{AB} < -10k_B T$, respectively. **b,f** Small-world parameters $\sigma_{sw}$ and $\omega_{sw}$ for condensates formed by YS sequences with varying hydrophobicity with energy threshold $E_{AB} < -k_B T$ and $E_{AB} < -10k_B T$, respectively. **d,j** Small-world parameters $\sigma_{sw}$ and $\omega_{sw}$ for condensates formed by YS sequences with $f_Y = 0.2$ and varying patterning with energy threshold $E_{AB} < -k_B T$ and $E_{AB} < -10k_B T$, respectively.

* These authors contributed equally

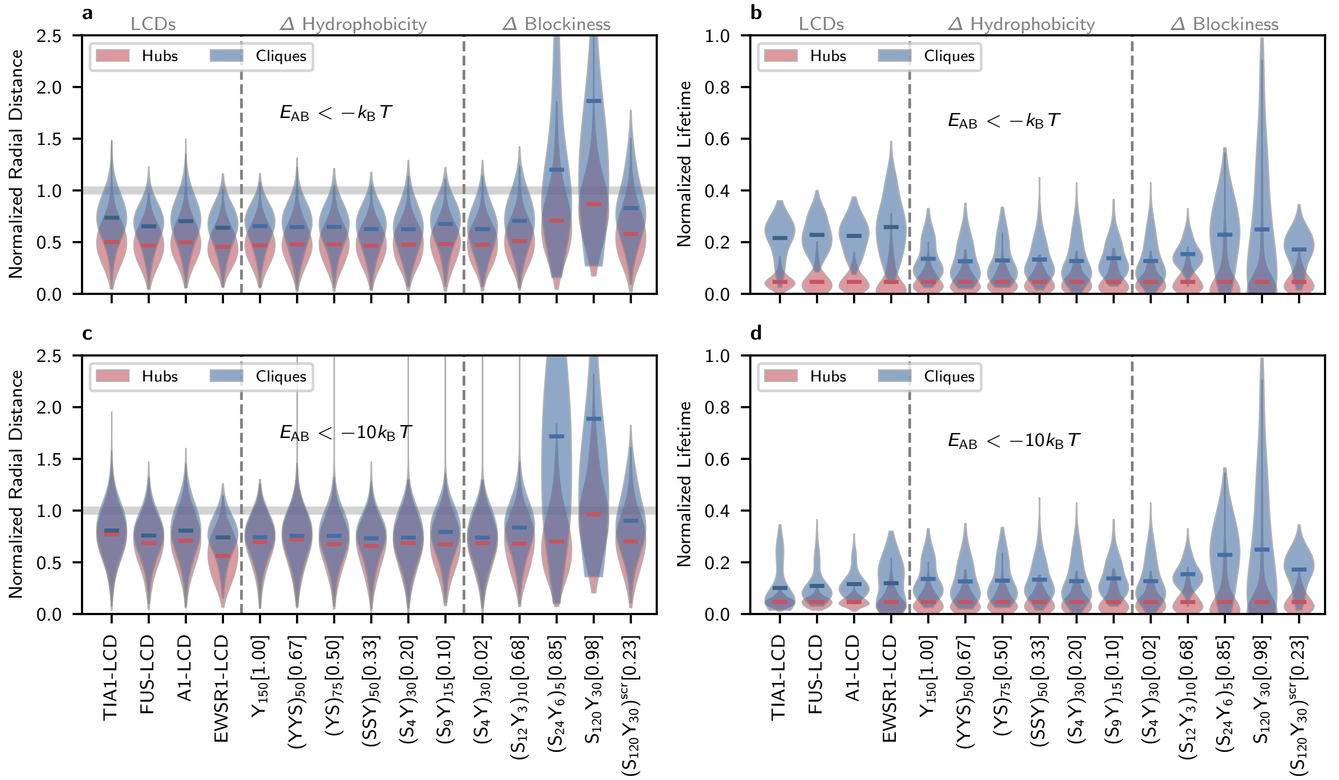


FIG. S2. **a,c** The distribution of hubs and cliques for all simulated sequences in terms of normalized radial distance from the condensate center of mass to interface with energy threshold $E_{AB} < -k_B T$ and $E_{AB} < -10k_B T$, respectively. Distances are normalized by the radial location of the interface in each simulation, overlaid in grey. YS sequences with varying hydrophobicity and conserved patterning are labeled with their hydrophobic fraction $f_Y$. YS sequences with varying patterning and conserved composition are labeled with their sequence blockiness $f_B$, described in the Methods. **b,d** Normalized lifetimes of hub molecules and clique molecules are shown for all simulated sequences with energy threshold $E_{AB} < -k_B T$ and $E_{AB} < -10k_B T$, respectively. Lifetimes are normalized by 200 ns, the length of the continuous trajectory sample used to generate these measurements. YS variants are labeled with $f_Y$ ($\Delta$ hydrophobicity) and $f_B$ ($\Delta$ blockiness) as described in (a,c).

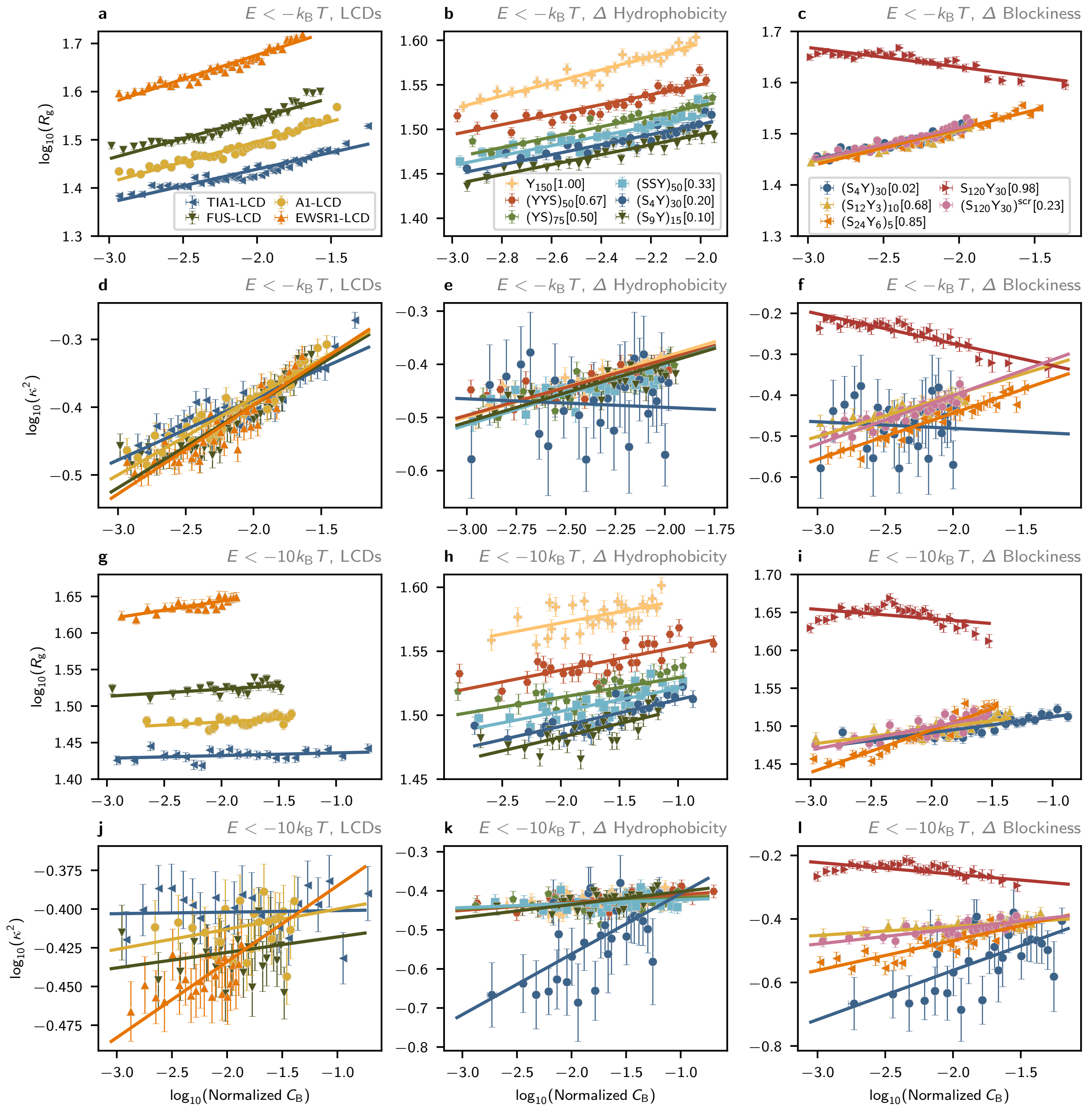


FIG. S3. **a, b, c, g, h, i** Possible power-law relations between the betweenness centrality $C_{\rm B}$ and radius of gyration $R_{\rm g}$, indicated by linear fits in $\log_{10}$–$\log_{10}$ space with energy threshold $E_{\rm AB} < -k_{\rm B}T$ and $E_{\rm AB} < -10k_{\rm B}T$, respectively.. **d, e, f, j, k, l** Possible power-law relations between the betweenness centrality $C_{\rm B}$ and relative shape anisotropy $\kappa^2$ with energy threshold $E_{\rm AB} < -k_{\rm B}T$ and $E_{\rm AB} < -10k_{\rm B}T$, respectively.

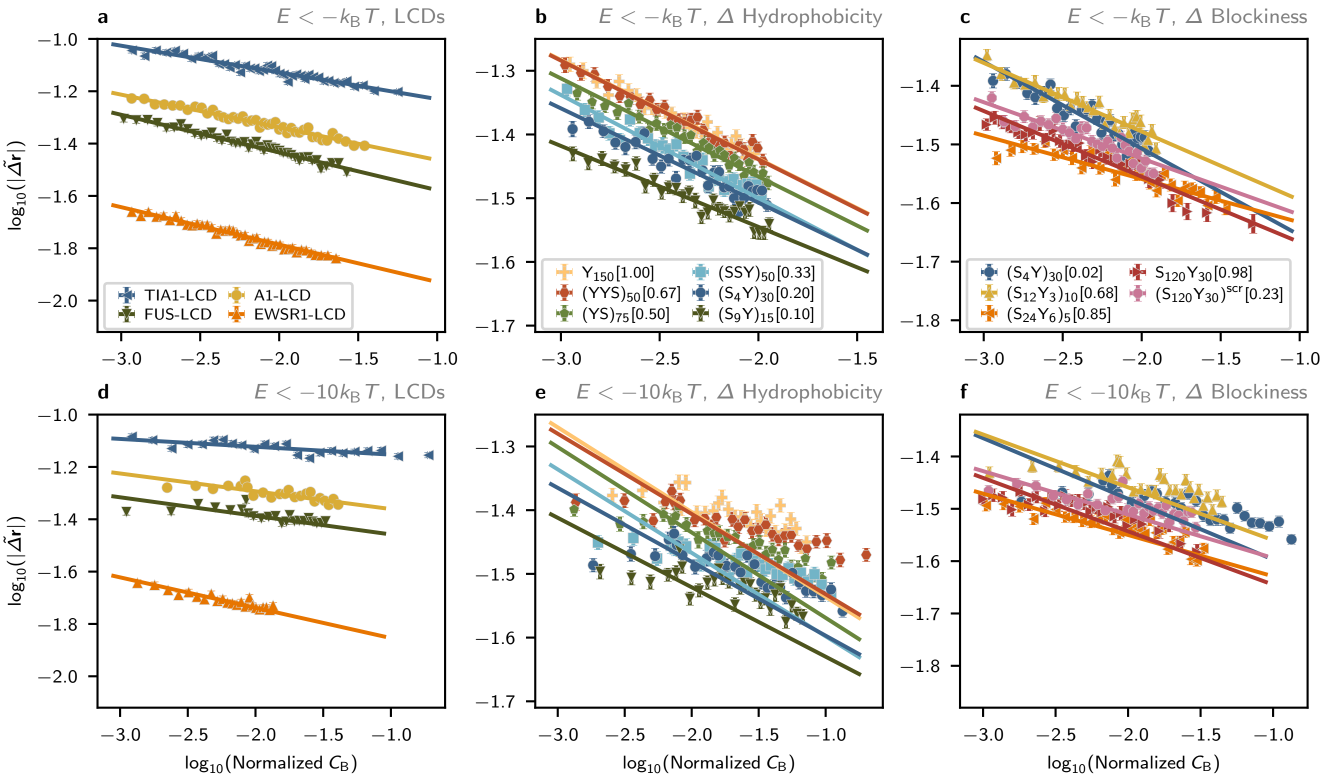


FIG. S4. **a, b, c** Normalized instantaneous displacements $|\tilde{\Delta}\mathbf{r}|$ of single chains in condensed phases exhibit strong negative correlations with node betweenness centrality $C_\mathrm{B}$ within molecular graphs with energy threshold $E_\mathrm{AB} < -k_\mathrm{B}T$. **d, e, f** Normalized instantaneous displacements $|\tilde{\Delta}\mathbf{r}|$ of single chains in condensed phases exhibit strong negative correlations with node betweenness centrality $C_\mathrm{B}$ within molecular graphs with energy threshold $E_\mathrm{AB} < -10k_\mathrm{B}T$.

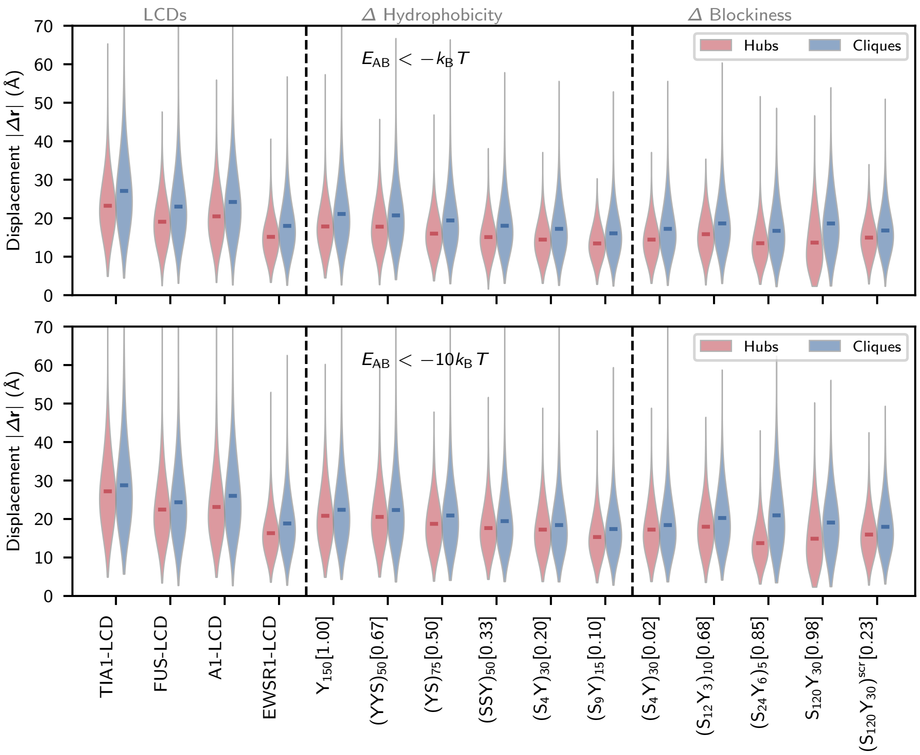


FIG. S5. **a, b** Displacements within consistent time intervals are compared for hub molecules and clique molecules with energy threshold $E_{AB} < -k_B T$ and $E_{AB} < -10k_B T$ (top and bottom, respectively).

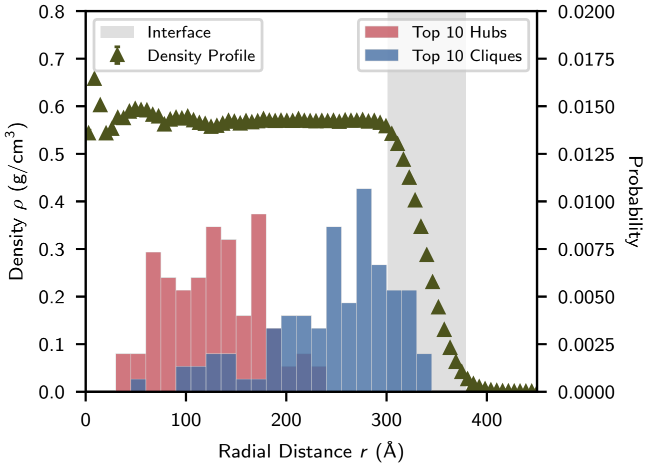


FIG. S6. Spatial distributions of hubs and cliques in a single-component FUS-LCD condensate containing 3375 chains at $T = 300$K. A radial mass density profile is shown in green triangles, and the phase interface is overlaid in grey. Consistent with results from simulations of smaller systems, clique molecules are closer to the condensate interface than hub molecules.

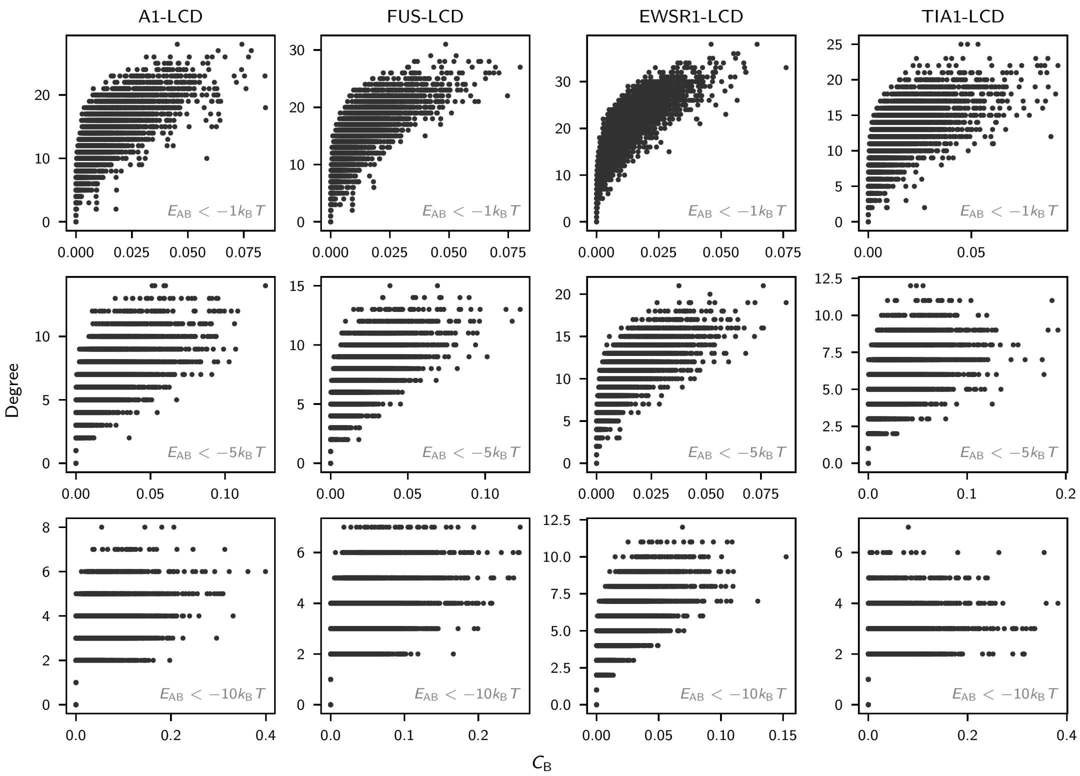


FIG. S7. Correlation between node degree and normalized betweenness centrality values $C_B$ in interaction networks constructed for LCD simulations at $T = 0.90T_c$. Graphs with edges assigned using the $E_{AB} < -1k_BT$ threshold are shown in the top row, $E_{AB} < -5k_BT$ in the center row, and $E_{AB} < -10k_BT$ in the bottom row. The positive correlation between node degree and betweenness centrality reflects the tendency of a node to connect more graph substructures when it connects many nodes, but the spread in the data indicate that a high node degree is not required for high betweenness centrality and vice versa.

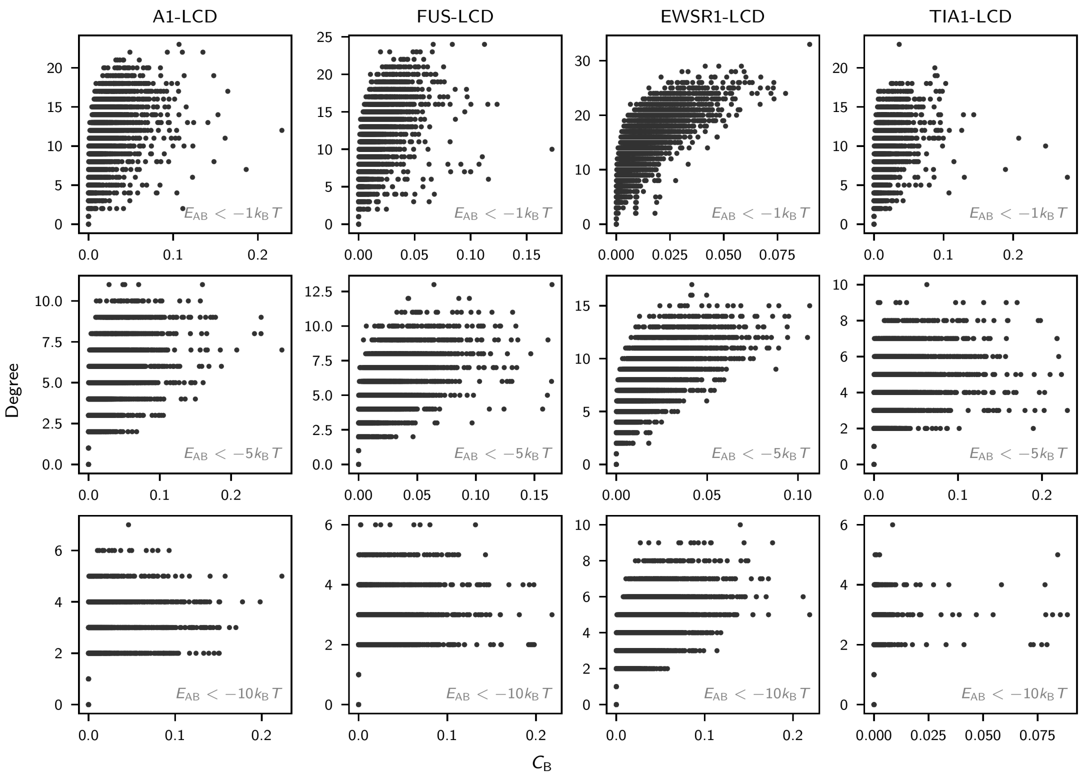


FIG. S8. Correlation between node degree and normalized betweenness centrality values $C_B$ in interaction networks constructed for LCD simulations at $T = 0.95T_c$. Graphs with edges assigned using the $E_{AB} < -1k_B T$ threshold are shown in the top row, $E_{AB} < -5k_B T$ in the center row, and $E_{AB} < -10k_B T$ in the bottom row.

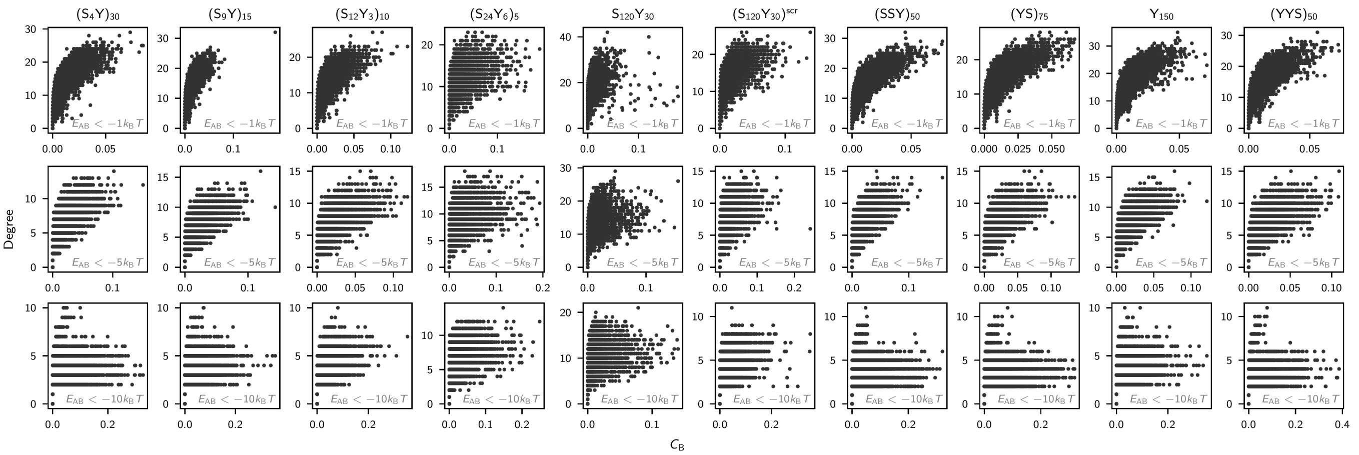

FIG. S9. Correlation between node degree and normalized betweenness centrality values $C_\mathrm{B}$ in interaction networks constructed for simulations of YS variants at $T = 0.90T_\mathrm{c}$. Graphs with edges assigned using the $E_\mathrm{AB} < -1k_\mathrm{B}T$ threshold are shown in the top row, $E_\mathrm{AB} < -5k_\mathrm{B}T$ in the center row, and $E_\mathrm{AB} < -10k_\mathrm{B}T$ in the bottom row. Though a positive trend is visible, the spread in the data indicate that high node degrees are not required for high betweenness centralities and vice versa.

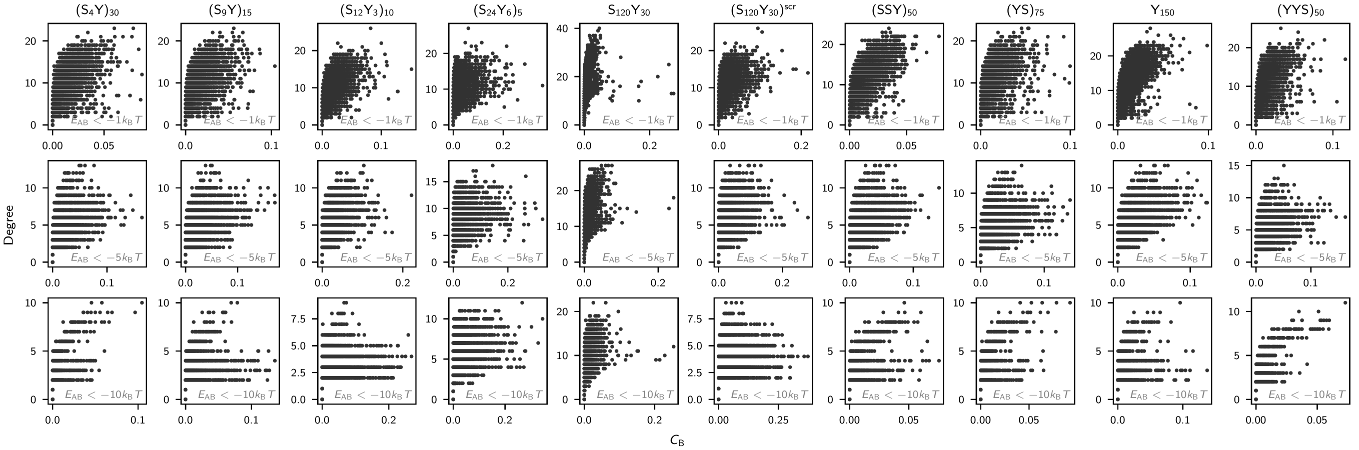

FIG. S10. Correlation between node degree and normalized betweenness centrality values $C_\mathrm{B}$ in interaction networks constructed for simulations of YS variants at $T = 0.95T_\mathrm{c}$. Graphs with edges assigned using the $E_\mathrm{AB} < -1k_\mathrm{B}T$ threshold are shown in the top row, $E_\mathrm{AB} < -5k_\mathrm{B}T$ in the center row, and $E_\mathrm{AB} < -10k_\mathrm{B}T$ in the bottom row.

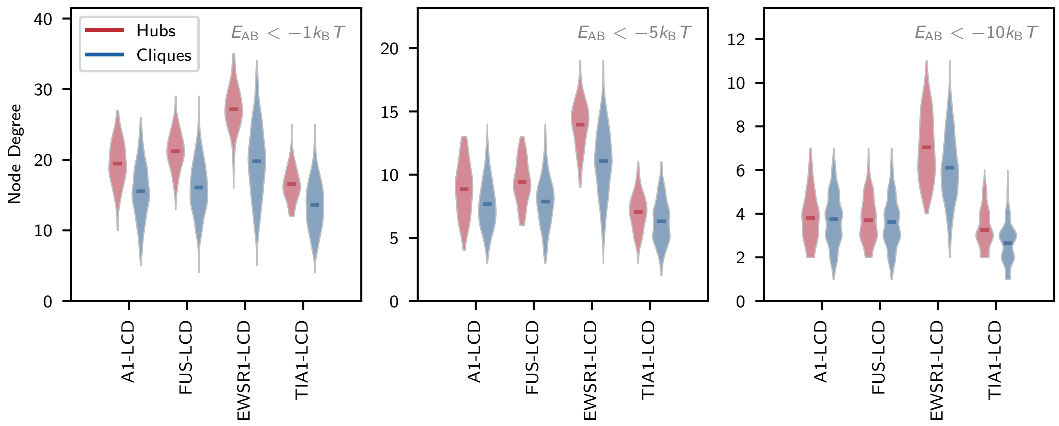


FIG. S11. Node degree distributions recorded for hub molecules (colored red) and clique molecules (colored blue), shown for simulations of LCD sequences at $T = 0.90T_c$. On average, hub molecules assume a higher degree (interact with more associative partners) than clique molecules, as expected. However, their overlapping distributions indicate that qualities of mesoscale organization, such as hublike or clique-like character, are inaccessible by analyzing degree alone.

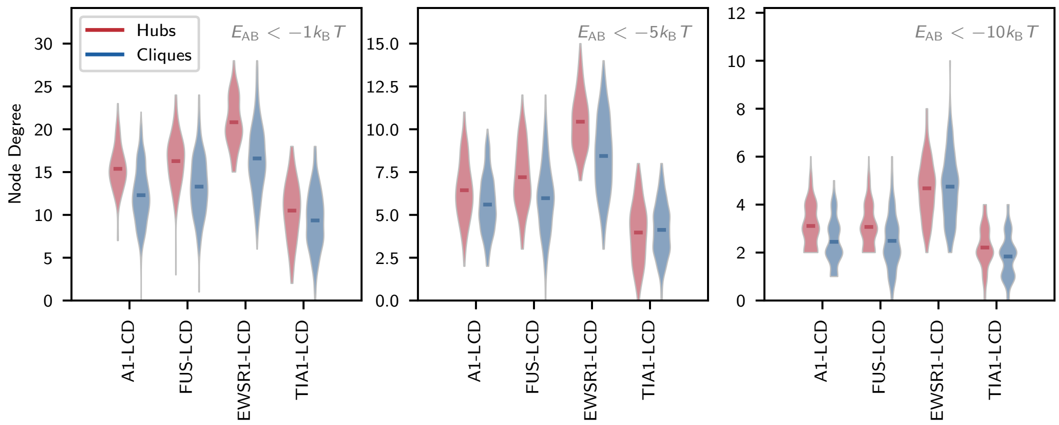


FIG. S12. Node degree distributions recorded for hub molecules (colored red) and clique molecules (colored blue), shown for simulations of LCD sequences at $T = 0.95T_c$.

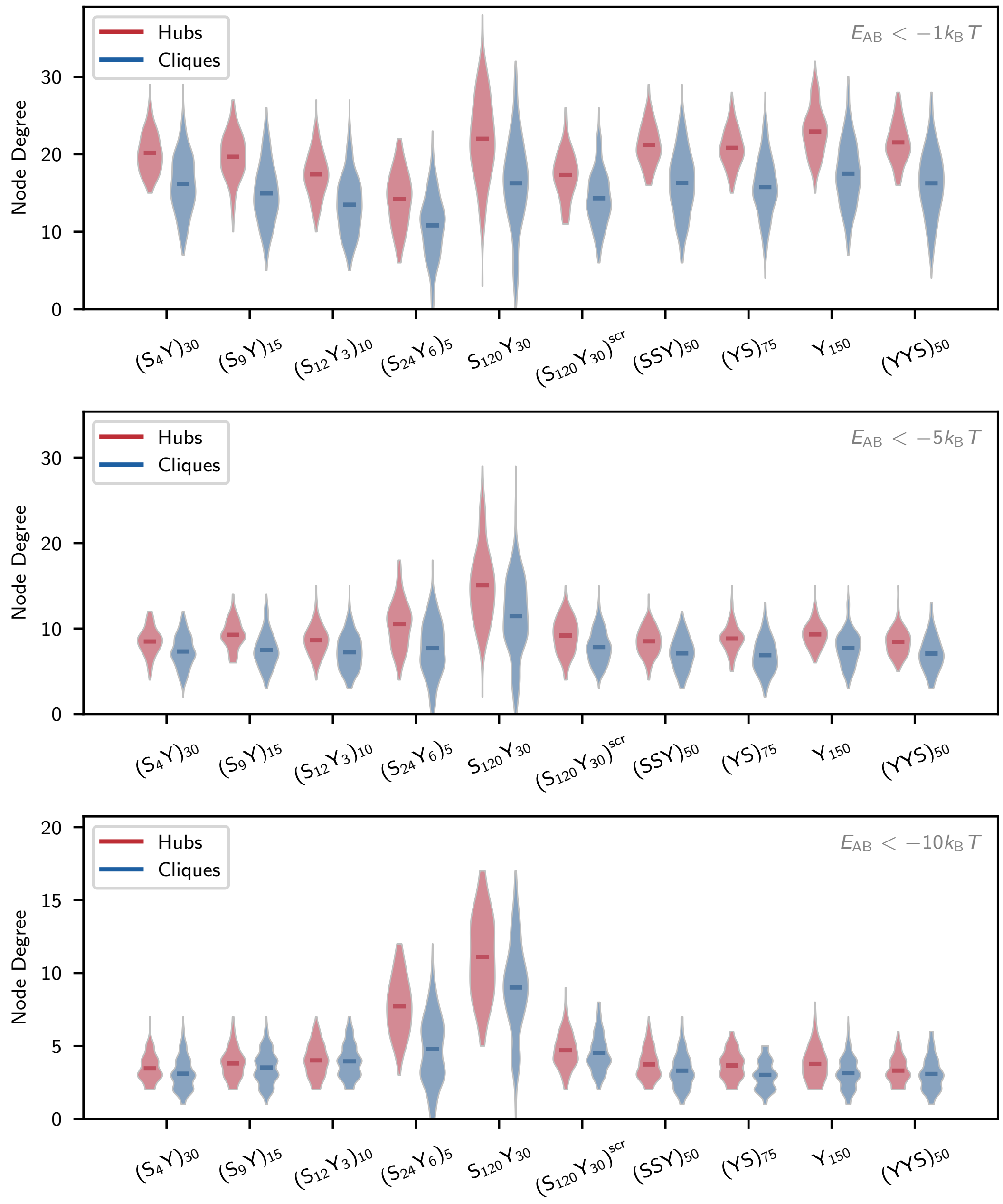


FIG. S13. Node degree distributions recorded for hub molecules (colored red) and clique molecules (colored blue), shown for simulations of YS variant sequences at $T = 0.90\,T_c$. On average, hub molecules assume a higher degree (interact with more associative partners) than clique molecules, as expected. However, their overlapping distributions indicate that qualities of mesoscale organization, such as hublike or clique-like character, are inaccessible by analyzing degree alone.

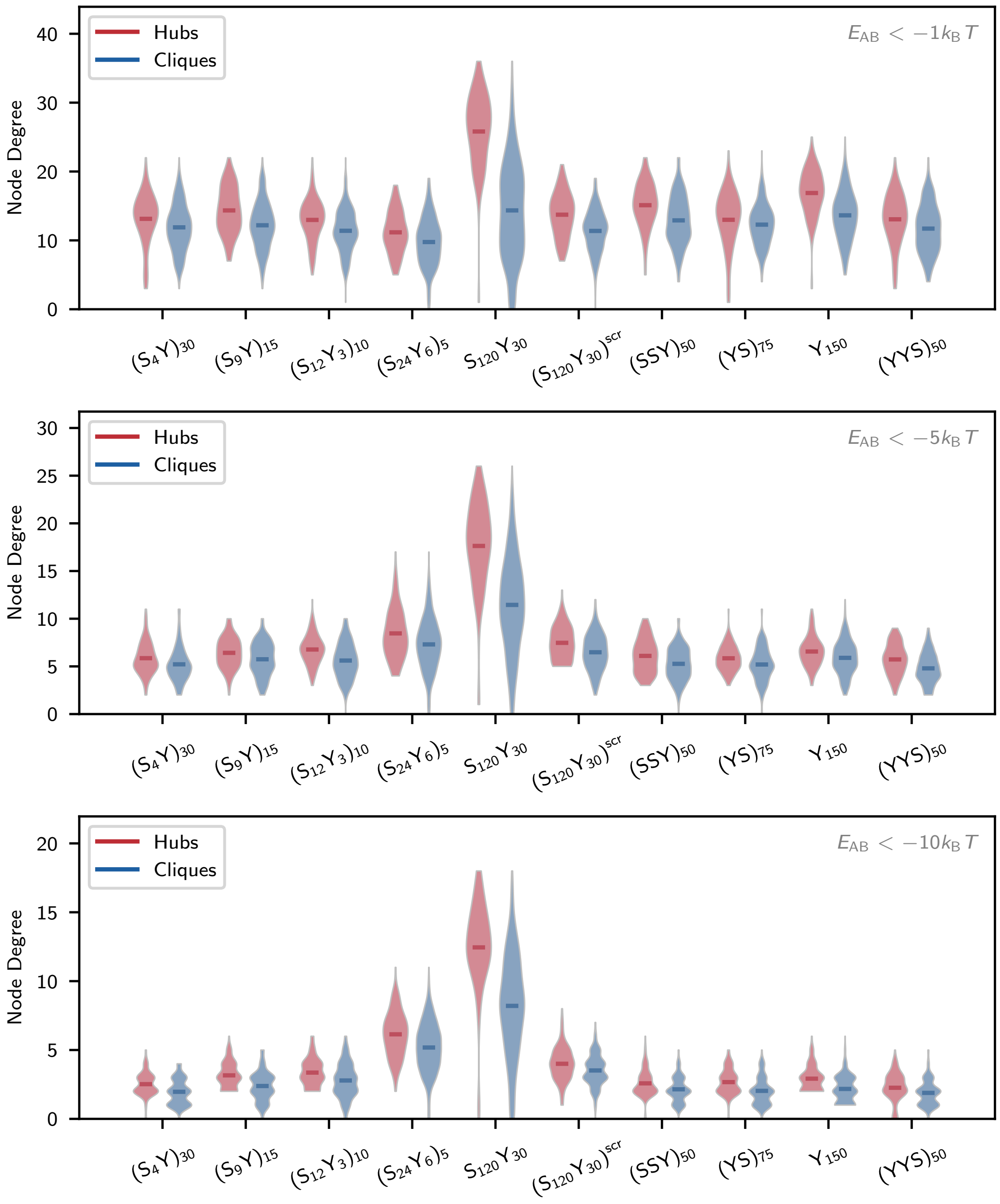


FIG. S14. Node degree distributions recorded for hub molecules (colored red) and clique molecules (colored blue), shown for simulations of YS variant sequences at $T = 0.90\,T_c$.

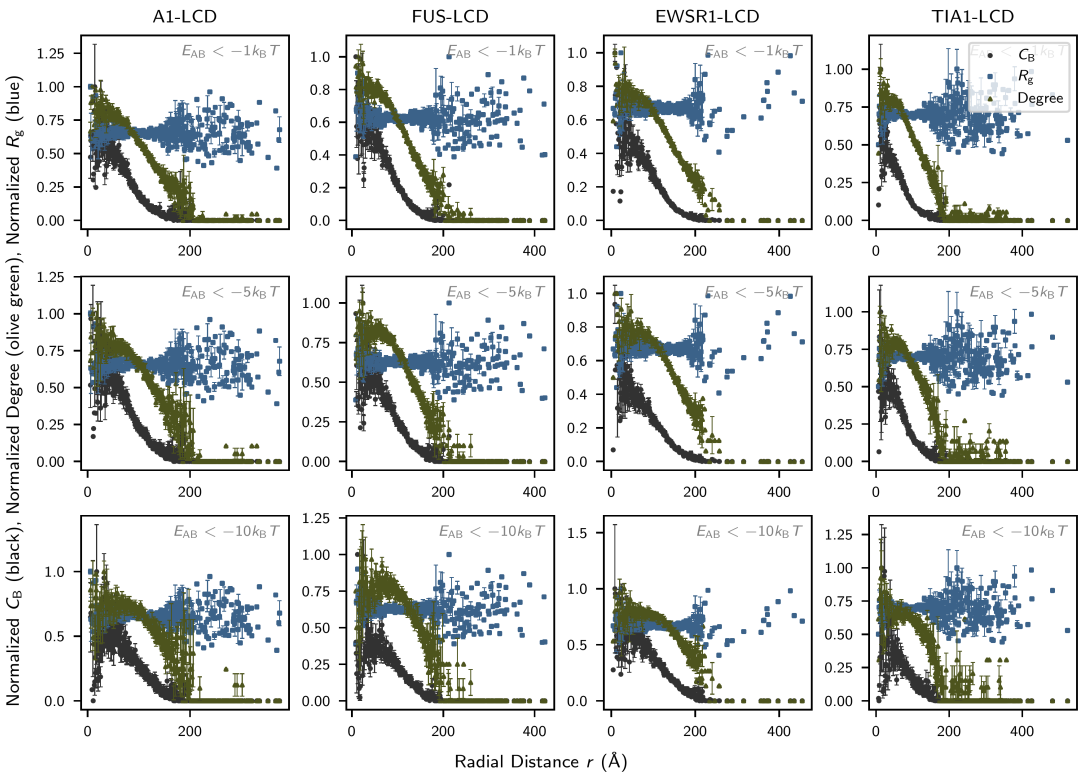


FIG. S15. Average dependence of node betweenness centrality (black circles), node degree (green triangles), and chain radius of gyration (blue squares) on the radial distance from the condensate center, shown for simulations of LCD sequences at $T = 0.90T_c$. Data for graphs with edges assigned using the $E_{AB} < -1k_BT$ threshold are shown in the top row, $E_{AB} < -5k_BT$ in the center row, and $E_{AB} < -10k_BT$ in the bottom row. $C_B$, degree, and $R_g$ (Å) are normalized by their maximal values in each subplot to aid visualization. Maximal values for each metric (i.e., inverse normalization coefficients) are presented in Table S1.

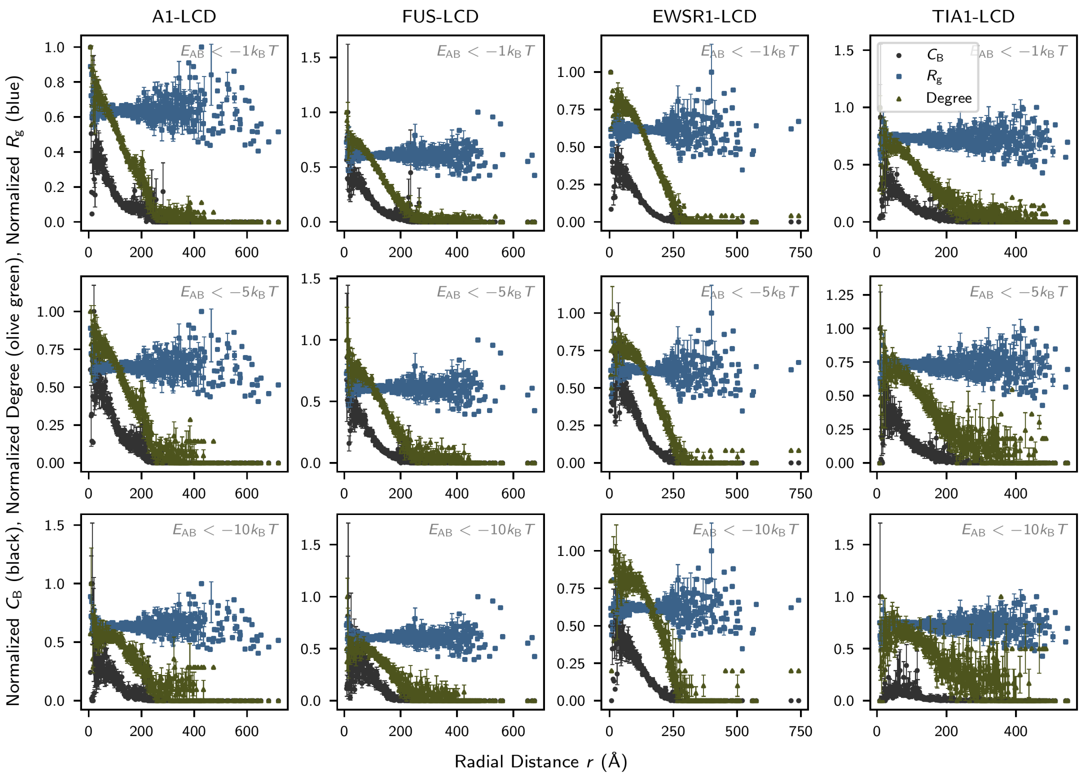


FIG. S16. Average dependence of node betweenness centrality (black circles), node degree (green triangles), and chain radius of gyration (blue squares) on the radial distance from the condensate center, shown for simulations of LCD sequences at $T = 0.95T_c$. Data for graphs with edges assigned using the $E_{AB} < -1k_BT$ threshold are shown in the top row, $E_{AB} < -5k_BT$ in the center row, and $E_{AB} < -10k_BT$ in the bottom row. Values are normalized by their maximal value in each subplot to aid visualization. Maximal values for each metric (i.e., inverse normalization coefficients) are presented in Table S1.

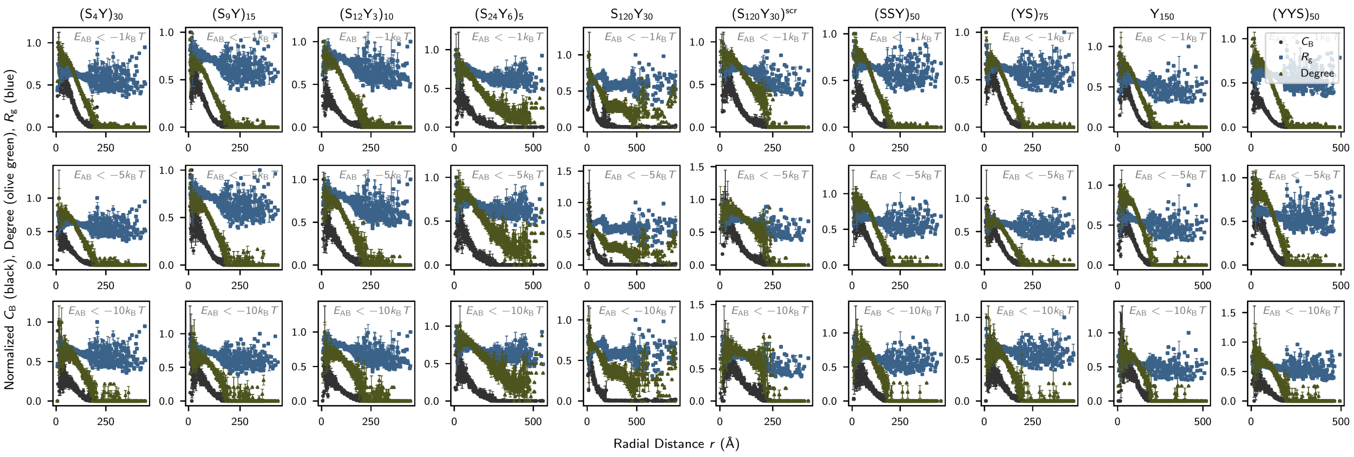


FIG. S17. Average dependence of node betweenness centrality (black circles), node degree (green triangles), and chain radius of gyration (blue squares) on the radial distance from the condensate center, shown for simulations of YS variants at $T = 0.90T_c$. Data for graphs with edges assigned using the $E_{AB} < -1k_BT$ threshold are shown in the top row, $E_{AB} < -5k_BT$ in the center row, and $E_{AB} < -10k_BT$ in the bottom row. Values are normalized by their maximal value in each subplot to aid visualization. Maximal values for each metric (i.e., inverse normalization coefficients) are presented in Table S2.

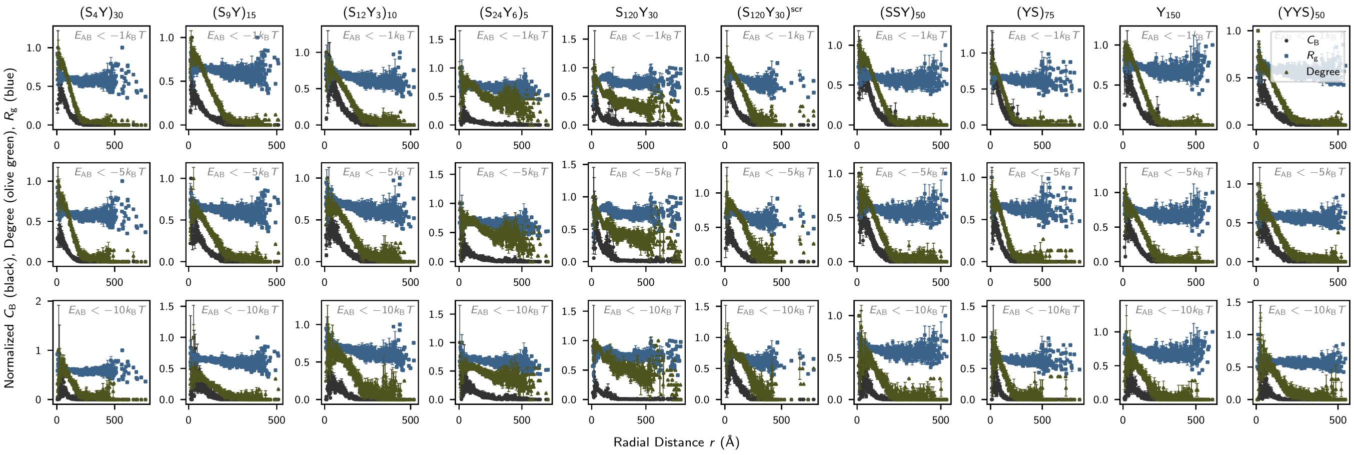


FIG. S18. Average dependence of node betweenness centrality (black circles), node degree (green triangles), and chain radius of gyration (blue squares) on the radial distance from the condensate center, shown for simulations of YS variants at $T = 0.95T_c$. Data for graphs with edges assigned using the $E_{AB} < -1k_BT$ threshold are shown in the top row, $E_{AB} < -5k_BT$ in the center row, and $E_{AB} < -10k_BT$ in the bottom row. Values are normalized by their maximal value in each subplot to aid visualization. Maximal values for each metric (i.e., inverse normalization coefficients) are presented in Table S3.

| Sequence | Temperature $/T_c$ | Energy Threshold $/-k_BT$ | Max $C_B$ | Max Degree | Max $R_g$ /Å |
|---|---|---|---|---|---|
| A1-LCD | 0.90 | 1 | 0.0419 | 21.0 | 45.8911 |
| A1-LCD | 0.90 | 5 | 0.0550 | 9.5 | 45.8911 |
| A1-LCD | 0.90 | 10 | 0.0992 | 4.0769 | 45.8911 |
| A1-LCD | 0.95 | 1 | 0.0709 | 18.0 | 47.0154 |
| A1-LCD | 0.95 | 5 | 0.0666 | 7.0 | 47.0154 |
| A1-LCD | 0.95 | 10 | 0.0416 | 3.5 | 47.0154 |
| FUS-LCD | 0.90 | 1 | 0.0420 | 23.3 | 53.8668 |
| FUS-LCD | 0.90 | 5 | 0.0566 | 10.0 | 53.8668 |
| FUS-LCD | 0.90 | 10 | 0.1214 | 4.0 | 53.8668 |
| FUS-LCD | 0.95 | 1 | 0.0599 | 20.0 | 55.7586 |
| FUS-LCD | 0.95 | 5 | 0.0823 | 8.0 | 55.7586 |
| FUS-LCD | 0.95 | 10 | 0.0391 | 4.0 | 55.7586 |
| EWSR1-LCD | 0.90 | 1 | 0.0446 | 32.0 | 66.4045 |
| EWSR1-LCD | 0.90 | 5 | 0.0614 | 16.0 | 66.4045 |
| EWSR1-LCD | 0.90 | 10 | 0.0574 | 7.5 | 66.4045 |
| EWSR1-LCD | 0.95 | 1 | 0.0720 | 24.0 | 73.2549 |
| EWSR1-LCD | 0.95 | 5 | 0.0586 | 12.0 | 73.2549 |
| EWSR1-LCD | 0.95 | 10 | 0.1088 | 5.0 | 73.2549 |
| TIA1-LCD | 0.90 | 1 | 0.0618 | 18.0 | 38.0820 |
| TIA1-LCD | 0.90 | 5 | 0.0785 | 7.3 | 38.0820 |
| TIA1-LCD | 0.90 | 10 | 0.0742 | 3.25 | 38.0820 |
| TIA1-LCD | 0.95 | 1 | 0.0510 | 14.0 | 35.9690 |
| TIA1-LCD | 0.95 | 5 | 0.0716 | 5.5 | 35.9690 |
| TIA1-LCD | 0.95 | 10 | 0.0057 | 2.0 | 35.9690 |

TABLE S1. Maximal measurement values for node betweenness centrality $C_B$ (unitless), node degree (unitless), and single-chain radius of gyration $R_g$ (Å) from simulations of LCD sequences. Tabulated values are used to normalize measurements of $C_B$, degree, and $R_g$ in visualizing their relationship to molecular distance from the condensate center of mass in Figs. S15 and S16.

| Sequence | Temperature $/T_c$ | Energy Threshold $/-k_BT$ | Max $C_B$ | Max Degree | Max $R_g$ /Å |
|---|---|---|---|---|---|
| $(S_4Y)_{30}$ | 0.90 | 1 | 0.0404 | 22.0 | 50.5730 |
| $(S_4Y)_{30}$ | 0.90 | 5 | 0.094 | 10.0 | 50.5730 |
| $(S_4Y)_{30}$ | 0.90 | 10 | 0.1672 | 4.6 | 50.5730 |
| $(S_9Y)_{15}$ | 0.90 | 1 | 0.0519 | 26.0 | 42.9463 |
| $(S_9Y)_{15}$ | 0.90 | 5 | 0.0810 | 12.0 | 42.9463 |
| $(S_9Y)_{15}$ | 0.90 | 10 | 0.1270 | 4.8 | 42.9463 |
| $(S_{12}Y_3)_{10}$ | 0.90 | 1 | 0.0777 | 19.0 | 39.9003 |
| $(S_{12}Y_3)_{10}$ | 0.90 | 5 | 0.0762 | 10.0 | 39.9003 |
| $(S_{12}Y_3)_{10}$ | 0.90 | 10 | 0.1622 | 5.0 | 39.9003 |
| $(S_{24}Y_6)_5$ | 0.90 | 1 | 0.0743 | 16.0 | 43.5978 |
| $(S_{24}Y_6)_5$ | 0.90 | 5 | 0.0770 | 11.0 | 43.5978 |
| $(S_{24}Y_6)_5$ | 0.90 | 10 | 0.0916 | 8.0 | 43.5978 |
| $S_{120}Y_{30}$ | 0.90 | 1 | 0.0335 | 28.0 | 64.8506 |
| $S_{120}Y_{30}$ | 0.90 | 5 | 0.0409 | 22.5 | 64.8506 |
| $S_{120}Y_{30}$ | 0.90 | 10 | 0.0476 | 15.0 | 64.8506 |
| $(S_{120}Y_{30})^{scr}$ | 0.90 | 1 | 0.0504 | 18.5 | 46.9385 |
| $(S_{120}Y_{30})^{scr}$ | 0.90 | 5 | 0.0534 | 9.2 | 46.9385 |
| $(S_{120}Y_{30})^{scr}$ | 0.90 | 10 | 0.0750 | 5.0 | 46.9385 |
| $(SSY)_{50}$ | 0.90 | 1 | 0.0601 | 24.0 | 50.2288 |
| $(SSY)_{50}$ | 0.90 | 5 | 0.0535 | 9.0 | 50.2288 |
| $(SSY)_{50}$ | 0.90 | 10 | 0.1317 | 3.8 | 50.2288 |
| $(YS)_{75}$ | 0.90 | 1 | 0.0427 | 26.0 | 53.5352 |
| $(YS)_{75}$ | 0.90 | 5 | 0.0641 | 12.0 | 53.5352 |
| $(YS)_{75}$ | 0.90 | 10 | 0.1508 | 4.8 | 53.5352 |
| $Y_{150}$ | 0.90 | 1 | 0.0401 | 28.0 | 68.4950 |
| $Y_{150}$ | 0.90 | 5 | 0.0545 | 10.0 | 68.4950 |
| $Y_{150}$ | 0.90 | 10 | 0.0976 | 4.5 | 68.4950 |
| $(YYS)_{50}$ | 0.90 | 1 | 0.0740 | 25.0 | 59.1013 |
| $(YYS)_{50}$ | 0.90 | 5 | 0.0601 | 9.1 | 59.1013 |
| $(YYS)_{50}$ | 0.90 | 10 | 0.1495 | 4.0 | 59.1013 |

TABLE S2. Maximal measurement values for node betweenness centrality $C_B$ (unitless), node degree (unitless), and single-chain radius of gyration $R_g$ (Å) from simulations of YS variant sequences at $T = 0.90T_c$. Tabulated values are used to normalize measurements of $C_B$, degree, and $R_g$ in visualizing their relationship to molecular distance from the condensate center of mass in Fig. S17.

| Sequence | Temperature $/T_c$ | Energy Threshold $/-k_BT$ | Max $C_B$ | Max Degree | Max $R_g$ /Å |
|---|---|---|---|---|---|
| $(S_4Y)_{30}$ | 0.95 | 1 | 0.0485 | 16.5 | 52.8052 |
| $(S_4Y)_{30}$ | 0.95 | 5 | 0.1055 | 7.0 | 52.8052 |
| $(S_4Y)_{30}$ | 0.95 | 10 | 0.0133 | 3.0 | 52.8052 |
| $(S_9Y)_{15}$ | 0.95 | 1 | 0.0734 | 17.0 | 48.0658 |
| $(S_9Y)_{15}$ | 0.95 | 5 | 0.0906 | 9.5 | 48.0658 |
| $(S_9Y)_{15}$ | 0.95 | 10 | 0.0295 | 6.0 | 48.0658 |
| $(S_{12}Y_3)_{10}$ | 0.95 | 1 | 0.0438 | 16.5 | 45.8733 |
| $(S_{12}Y_3)_{10}$ | 0.95 | 5 | 0.0793 | 9.0 | 45.8733 |
| $(S_{12}Y_3)_{10}$ | 0.95 | 10 | 0.1414 | 4.0 | 45.8733 |
| $(S_{24}Y_6)_5$ | 0.95 | 1 | 0.1861 | 13.0 | 43.9439 |
| $(S_{24}Y_6)_5$ | 0.95 | 5 | 0.1470 | 10.0 | 43.9439 |
| $(S_{24}Y_6)_5$ | 0.95 | 10 | 0.1270 | 9.0 | 43.9439 |
| $S_{120}Y_{30}$ | 0.95 | 1 | 0.0323 | 27.3 | 56.5004 |
| $S_{120}Y_{30}$ | 0.95 | 5 | 0.0234 | 19.0 | 56.5004 |
| $S_{120}Y_{30}$ | 0.95 | 10 | 0.0652 | 12.0 | 56.5004 |
| $(S_{120}Y_{30})^{scr}$ | 0.95 | 1 | 0.0582 | 14.5 | 44.7216 |
| $(S_{120}Y_{30})^{scr}$ | 0.95 | 5 | 0.0926 | 7.7 | 44.7216 |
| $(S_{120}Y_{30})^{scr}$ | 0.95 | 10 | 0.0941 | 4.0 | 44.7216 |
| $(SSY)_{50}$ | 0.95 | 1 | 0.0342 | 18.0 | 54.5406 |
| $(SSY)_{50}$ | 0.95 | 5 | 0.0661 | 8.2 | 54.5406 |
| $(SSY)_{50}$ | 0.95 | 10 | 0.0228 | 4.0 | 54.5406 |
| $(YS)_{75}$ | 0.95 | 1 | 0.0293 | 17.5 | 50.4438 |
| $(YS)_{75}$ | 0.95 | 5 | 0.0623 | 7.5 | 50.4438 |
| $(YS)_{75}$ | 0.95 | 10 | 0.0122 | 2.7 | 50.4438 |
| $Y_{150}$ | 0.95 | 1 | 0.0352 | 18.6 | 51.9253 |
| $Y_{150}$ | 0.95 | 5 | 0.0640 | 8.0 | 51.9253 |
| $Y_{150}$ | 0.95 | 10 | 0.0279 | 3.6 | 51.9253 |
| $(YYS)_{50}$ | 0.95 | 1 | 0.0544 | 22.0 | 59.4124 |
| $(YYS)_{50}$ | 0.95 | 5 | 0.0682 | 8.0 | 59.4124 |
| $(YYS)_{50}$ | 0.95 | 10 | 0.0150 | 3.2 | 59.4124 |

TABLE S3. Maximal measurement values for node betweenness centrality $C_B$ (unitless), node degree (unitless), and single-chain radius of gyration $R_g$ (Å) from simulations of YS variant sequences at $T = 0.95T_c$. Tabulated values are used to normalize measurements of $C_B$, degree, and $R_g$ in visualizing their relationship to molecular distance from the condensate center of mass in Fig. S18.
